# High-resolution mapping of human RNA polymerase III reveals transcription termination as a rate-limiting step

**DOI:** 10.1101/2025.10.17.683076

**Authors:** Jan Mikołajczyk, Ameen Kollaran, Jonas Weidenhausen, Christoph W. Müller, Tomasz W. Turowski

## Abstract

Transfer RNA (tRNA) molecules play a central role in the flow of genetic information, translating nucleic acid sequences into the functional protein portfolio of an organism. Despite the high abundance of tRNAs in cells, studying their biology remains challenging due to the repetitive nature of genomic sequences and the numerous modifications of mature molecules. tRNA genes (tDNAs) are transcribed exclusively by RNA polymerase III (RNAPIII), which has been recently linked to rare genetic disorders but also longevity.

Here, we present the first mapping of actively transcribing RNAPIII in the human, using K562 cell line and UV-crosslinking followed by stringent purification. Our data reveals high variability in expression across tDNAs, common occurrences of transcriptional read-through, and unique transcription dynamics across transcription units, with the lowest kinetics associated with transcription termination. Further analysis revealed that release of the nascent transcript is a critical step in tRNA transcription. Unexpectedly, shorter terminators promote more efficient termination, which becomes the rate-limiting step for human RNAPIII transcription of highly expressed tDNAs.

Our dataset provides insights into actively transcribing RNAPIII dynamics in human cells, allowing for a comprehensive analysis of genomic loci transcribed by RNAPIII. Our high resolution data and resulting kinetic information reveal that human RNAPIII contradicts the paradigm that initial steps of transcription are the main determinants of its output. This is significant as RNAPIII is commonly used to transcribe synthetic RNA constructs, such as short hairpin RNAs or guide RNAs used in gene editing technologies. With our study we offer a functional framework for the analysis of RNAPIII activity, addressing the need for comprehensive understanding of RNAPIII transcription fueled by a growing number of described genetic disorders caused by mutations in this enzyme.

## INTRODUCTION

The human genome spans 3.1 billion base pairs (3.1 Gb), but only a fraction is actively transcribed at any given time. The majority of expressed genes are transcribed by RNA polymerase II (RNAPII), which synthesizes all protein-coding genes but also a wide array of non-coding RNAs. Protein-coding regions account for only 1–2% of the human genome, the presence of introns and untranslated regions expands protein-coding genes to cover approximately 40% of genomic DNA. In contrast, RNA polymerases I (RNAPI) and III (RNAPIII) are responsible for transcribing a relatively small portion of the genome, but collectively account for over 95% of RNA in a cell. RNAPI transcribes ribosomal DNA (rDNA) from three clusters in the human genome, while RNAPIII produces highly abundant, short non-coding RNAs, including several hundred tRNAs, U6 snRNA, 7SL RNA, 5S rRNA, and others [reviewed in ^1^]. RNAPIII also transcribes a subset of retrotransposons, which belong to genomic elements that collectively constitute up to ∼50% of the human genome^2^. Therefore, understanding RNAPIII is essential for a comprehensive view of human transcription.

Eukaryotic transcription is regulated at multiple levels, including chromatin structure and accessibility, transcription factor binding, initiation, and elongation. Until recently, transcription termination was not considered a major determinant of transcription output. However, inefficient termination can increase cellular burden and disrupt adjacent transcription units^3^.

Although tRNAs are often regarded as constitutively expressed housekeeping molecules, their transcription by RNAPIII is modulated by chromatin accessibility and histone modifications, which provide gene-specific regulation responsive to environmental and developmental cues^4–7^. Regulation of tDNA expression through selective activation and repression has long been proposed, yet the molecular mechanisms governing RNAPIII regulation remain poorly understood. In yeast, tRNA copy number correlates with the abundance of mature tRNAs, and general repression by the conserved Maf1 protein adjusts expression in response to environmental signals^8,9^. However, not all tDNAs respond equally to this regulation in yeast or mouse, suggesting the existence of a subset of constitutively active tRNA housekeeping genes^10,11^. While in yeast tDNAs are dispersed throughout the genome with no clear pattern, in humans, many tRNA genes are organized in linear genomic clusters.

Unlike RNAPI and RNAPII, which initiate transcription from upstream promoters, most RNAPIII promoters are located within the genes themselves. RNAPIII utilizes three classes of promoters; class II promoters are characteristic of the 70–80 bp tDNAs, with A- and B-box elements forming an internal promoter [reviewed in ^12^]. These elements are recognized by the τA and τB subcomplexes of the TFIIIC transcription factor^13^. TFIIIC recruits TFIIIB, and together they enable RNAPIII loading and initiation. However, RNAPIII elongation kinetics remain poorly understood, largely due to the short length of transcription units and the high abundance of mature transcripts, both of which pose significant technical challenges.

The precision of RNAPIII transcription initiation remains unclear. RNAPI begins transcription at very specific start sites, while RNAPII initiates across a wider area. Its transcription start sites (TSSs) can span several nucleotides and show different patterns of direction—ranging from completely bidirectional to unidirectional^14–16^. Both RNAPI and RNAPII display variable elongation kinetics, with slower progression near gene starts—potentially explained by gradual torsional entrainment mechanism^17^. In higher eukaryotes, the initial kinetics of RNAPII are tightly regulated by promoter-proximal pausing^18^.

A distinct feature of RNAPIII transcription is its termination mechanism, which involves recognition of a short stretch of thymidines located in the 3’end of the tDNA. This process is prone to failure, resulting in transcriptional read-through^11,19^. Recent findings suggest that sequences on the non-template strand, as well as the Sen1 protein, contribute to RNAPIII termination in yeast^20^. To date, human RNAPIII activity has primarily been studied using chromatin immunoprecipitation (ChIP) approaches, which reveal polymerase occupancy^10,21^, or metabolic labeling techniques such as PRO-seq, which measures global transcription activity^22,23^.

Here, we report the first transcriptome-wide mapping of actively transcribing RNAPIII in human cells. Our approach identifies which tDNAs are actively transcribed and delineates initiation and termination sites of RNAPIII at nucleotide resolution. The resolution of our data, obtained using a modified crosslinking and analysis of cDNA (CRAC) protocol, allows for kinetic analysis of transcription dynamics and reveals that transcription termination is the rate-limiting step in high-turnover tRNA transcription. Using reporter constructs, we demonstrate that a terminator consisting of four thymidines (4T) yields the highest levels of pre-tRNA by efficiently releasing nascent transcripts.

## RESULTS

### Mapping nascent transcripts of RNAPIII identifies transcribed tDNAs with extended 3′ ends and transcription of retrotransposable elements

To map actively transcribing RNAPIII in human cells, we employed the CRAC method, which we previously applied to study RNAPI and RNAPIII in yeast^12,15^. We engineered a K562 suspension cell line expressing the largest RNAPIII subunit, RPC1, tagged at the C-terminus with an 8×His-FLAG (HF) epitope. Cells were irradiated with UV light at 254 nm, resulting in covalent crosslinks between RNA and proteins. Subsequently the target protein, RPC1-HF, is purified with its attached interacting RNAs. A major strength of the CRAC approach lies in its high specificity, achieved through tandem affinity purification including stringent, denaturing conditions (6 M guanidinium hydrochloride) (Fig. 1A), enabling confident detection of abundant transcripts such as rRNAs and tRNAs.

**Figure 1.**
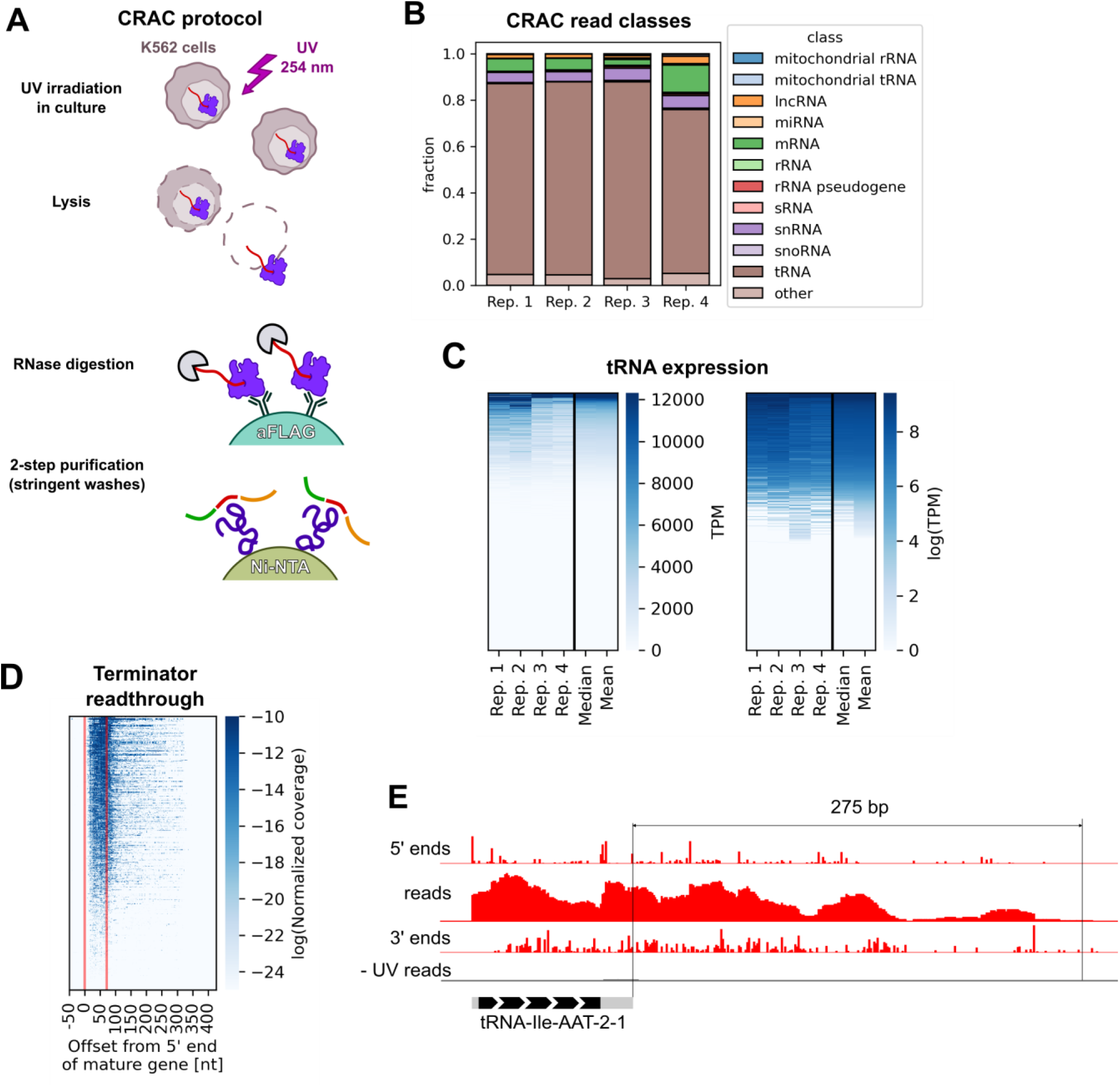
CRAC maps actively transcribing RNAPIII with nucleotide resolution. A: Outline of the CRAC protocol. RNA is crosslinked to proteins with 254 nm UV irradiation. After lysis, tagged protein is purified using affinity purification and washed in denaturing conditions. Library adapters (green and orange) are ligated on beads and the library is released by protein K digestion before reverse transcription and PCR amplification. B: Overview of RNAPIII CRAC reads mapping to different RNA classes. C: Heatmaps of tDNA expression. Biological replicates (n=4), as well as the mean and median are shown (left). Same data after logarithmic transformation reveals silent tDNAs (right). D: Heatmap showing profiles of CRAC read 3’ end coverage across tDNAs. Mature tRNA region is marked with red lines. E: An example CRAC profile of a tDNA (tRNA-Ile-AAT-2-1) with marked tDNA and 275 bp terminator readthrough.

To map the very 3′ ends of nascent transcripts, we used a modified version of the CRAC protocol that omits phosphatase treatment, allowing adaptor ligation only to 3′-OH ends. Additionally, data preprocessing steps included adaptor trimming and filtering for adaptor-containing reads to ensure accurate 3′ end assignment (Fig. S1A). The transcription start sites (TSSs) were inferred from length-filtered 5′ ends of mapped reads.

The dataset was highly reproducible across four independent biological replicates (Spearman’s R > 0.91; Fig. S1B) and showed strong enrichment for canonical RNAPIII transcript classes, with the majority of reads mapping to tDNAs (Fig. 1B). To quantify RNAPIII engagement at individual loci, we assigned reads to genomic features, providing a locus-specific profile of transcriptionally active RNAPIII—serving as a proxy for nascent transcript expression levels (see Limitations of the Study). Given the high sequence similarity among tDNAs, we applied two independent quantification strategies to ensure the robustness of our expression estimates (Fig. S1C-E, see Methods for details).

CRAC coverage across tDNAs revealed that ∼1/3 of predicted tDNAs are actively transcribed at relatively high levels, while nearly 50% appear transcriptionally silent in K562 cells (Fig. 1C). Importantly, CRAC reflects RNAPIII engaged in active transcription, offering a more precise snapshot of gene activity than occupancy-based methods such as ChIP. In addition to tRNAs, we detected, as expected, signals for other highly expressed non-coding RNAs (e.g., 7SL, 5S rRNA, U6 snRNA). RNAPIII activity at the neuron-specific *BC200* transcript was not detected, consistent with the cell-type specificity of its expression (Fig. S1K).^24^

RNAPIII was previously found to transcribe repetitive genomic elements such as Alu and B2 SINE^25^. Therefore, we examined RNAPIII activity at these elements in our data. CRAC reads were mapped to a reference containing predictions from the RetroSeeker pipeline^26^ (see Methods for details). Depending on quantification strategy, retrotransposon-derived transcripts accounted for ∼2–6% of total RNAPIII reads (Fig. S1H-I). Many of these loci include functional RNAPIII promoter elements (Fig. S1J), and several unknown RNAPIII transcripts were identified (Fig. S1L, Table S2).

We observed common RNAPIII readthrough at tDNAs (Fig. 1D, S1F), reminiscent of what we previously described in *S. cerevisiae*. Unexpectedly, in some cases, transcription extended several hundred base pairs beyond the primary poly(T) terminator, passing multiple downstream terminators before successful termination (Fig. 1E). A systematic analysis revealed that readthrough intensity is weakly anticorrelated with terminator strength (Fig. S1G). However, the rules governing tRNA termination readthrough remain unknown. These findings indicate that RNAPIII termination events in humans are often inefficient.

### Transcriptionally active tDNA are dependent on favourable chromatin state, but not higher order topological structures

The abundance of different tRNA isodecoders has usually been attributed to the number of corresponding tDNA copies in the genome. Therefore, we decided to investigate why the substantial fraction of tDNAs is transcriptionally silent (Fig. 1C), and what constitutes the regulatory mechanisms underlying this phenomenon. Chromatin accessibility, governed by protein factors and epigenetic modifications, is a key determinant of gene expression. To explore this, we analysed publicly available ChIP-seq datasets from the ENCODE project to assess the chromatin landscape surrounding tDNAs^24^ (for complete list of used datasets see Table S3). This revealed that RNAPIII activity correlates with specific chromatin marks such as H3K4me2, EP300, c-Jun and others (Fig. 2A and S2). RNAPIII activity revealed clear differences in chromatin status when compared to RNAPII and III occupancy measured with ChIP. Surprisingly, RNAPIII occupancy correlated less strongly with translationally active RNAPIII coverage than the aforementioned chromatin features, indicating differences between RNAPIII association with tDNA (occupancy) and transcriptional activity (nascent transcripts).

**Figure 2.**
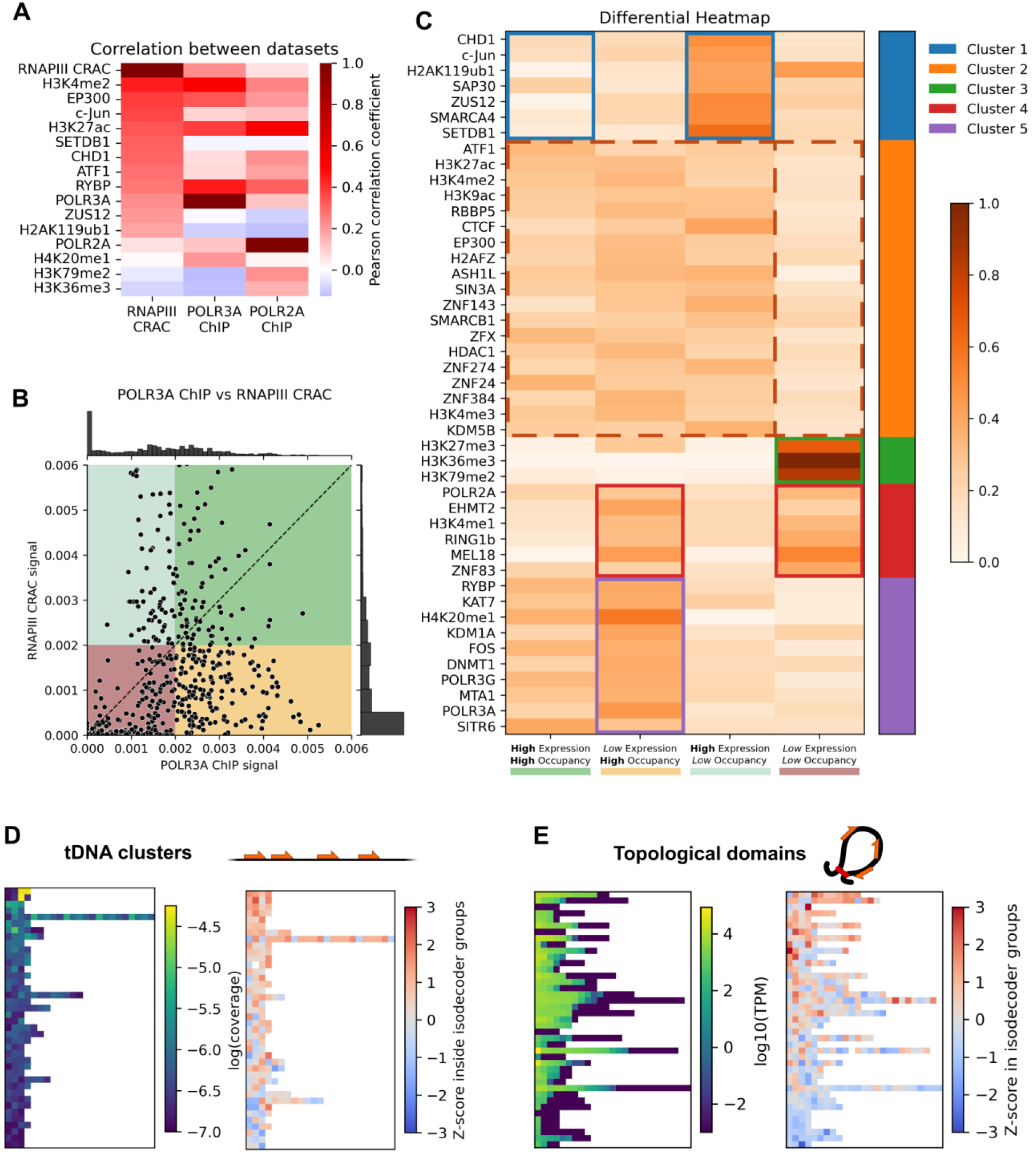
Chromatin-associated factors and epigenetic marks, but not topological conformation, are high-level tDNA expression regulators. A: Heatmap of Pearson correlation coefficients between RNAPIII CRAC data and ENCODE ChIP-seq datasets of selected chromatin-associated factors and epigenetic marks. B: Scatterplot of RNAPIII CRAC signal versus POLR3A ChIP-seq signal, plotted individually for every tDNA. C: Differential heatmap of chromatin-associated factor and mark binding. tDNAs were divided into 4 groups (columns) based on RNAPIII occupancy, inferred from POLR3A ChIP-seq, and expression levels, inferred from RNAPIII CRAC. Factors and marks were than clustered according to ChIP-seq signal levels across these groups. D: Expression relationships in linear tDNA clusters. Each heatmap row represents a cluster of tDNAs with individual genes spaced at most 5 kbp apart. Clusters containing less than three genes are not shown. E: tDNA expression in topological domains. Each heatmap row represents a cluster of tDNA located in a given topological domain determined by Hi-C data.

To systematically examine these differences, we categorized tDNAs into four groups based on their CRAC (expression) and ChIP (occupancy) signals: high expression and occupancy (green), high expression but low occupancy (sea green), low expression but high occupancy (yellow) and low occupancy and expression (red) (Fig. 2B). This revealed several clusters of chromatin factors presented in Fig. 2C: cluster 1 (blue) which is characteristic for genes with high expression but lower occupancy (i.e. c-Jun and ZUS12). Cluster 2 (orange) containing factors clearly depleted for low RNAPIII activity and occupancy. We envisage these factors play a critical role in establishing functional RNAPIII chromatin. An opposite function could be attributed to cluster 3 (green), which contains three types of H3 methylations, characteristic for RNAPIII absence.

Interestingly, cluster 4 (red) includes chromatin where RNAPIII is present but is not active. This could reflect inhibitory proteins and marks, in line with presence of RNAPII. Finally, cluster 5 (purple) represents chromatin status associated with RNAPIII occupancy, but not necessarily activity.We also generated metagene profiles of ChIP-seq signals for each group from Fig. 2B and calculated enrichment or depletion relative to the global tDNA set (Fig. S3), revealing differences in chromatin status at tDNA and surroundings. These data suggest that active transcription by RNAPIII is strongly influenced by chromatin state, particularly through mechanisms that modulate accessibility during initiation.

Given that many tDNAs are organized into linear clusters within the genome, we next asked whether such clustering contributes to co-regulation. To address this, we quantified CRAC signals as absolute read count (RNAPIII engagement, Fig. 2D, left panel) and as the percentage of total RNAPIII engagement assigned to each isodecoder (Z-score of isodecoder expression, Fig. 2D, right panel). The latter metric was designed to control for variability introduced by differences in A and B box strength within internal promoter elements (see Methods for details). This analysis revealed most (85 out of 114) clusters contain tDNAs with a distribution of expression levels similar to the global distribution for all tDNAs, and as such are unlikely to be regulated as units (Fig. 2B and S4A, Kolmogorov-Smirnov test p ≥ 0.05).

Finally, we investigated whether higher-order chromatin conformation might play a role in regulating tDNA expression. Using Hi-C data^27^, we identified topologically associating domains (TADs) and grouped tDNA accordingly. We again used two measures of expression, as in the analysis of linear clusters (Fig. 2E and S4B). Most domains contained tDNAs with a broad range of expression levels (Fig. 2E). To determine whether the expression distribution within each domain differed from the global distribution across all tDNAs, we applied the Kolmogorov-Smirnov test. A non-significant result (p ≥ 0.05) indicated that expression levels within a domain did not deviate from the global profile. This analysis showed that only 3 to 6 domains (depending on the window size used for determining TAD boundaries) exhibited significantly different tDNA expression level distributions. Thus, we conclude that while silent tDNAs show a slight predisposition to cluster spatially, genome topology is not a major determinant of RNAPIII-mediated tDNA expression at genome-wide scale.

### RNAPIII CRAC identifies unidirectional but sparse transcription initiation sites

When gene-encoding DNA becomes accessible, transcription initiation marks a critical step in RNA synthesis, defining the precise 5′ end of the nascent transcript—known as the transcription start site (TSS). While RNAPII exhibits dispersed and often bidirectional TSSs, RNAPI initiates transcription at a single, sharply defined nucleotide in a strictly unidirectional manner. This prompted us to investigate the nature of RNAPIII TSSs in human cells. To do so, we analysed peaks corresponding to the 5′ ends of CRAC reads. To minimize potential artifacts introduced by RNase digestion during library preparation, we limited the analysis to reads shorter than 20 nucleotides. Visual inspection of the 5′ leaders revealed two distinct classes of TSSs: dominant, primary TSSs (strongest) and weaker, upstream secondary TSSs (upstream) (Fig. 3A). These were classified using unbiased computational algorithms (see Methods). This allowed us to annotate human tDNA TSSs and revealed that human RNAPIII, like other polymerases, prefers to initiate transcription with purine residues (A or G) (Fig. 3B–C).

**Figure 3.**
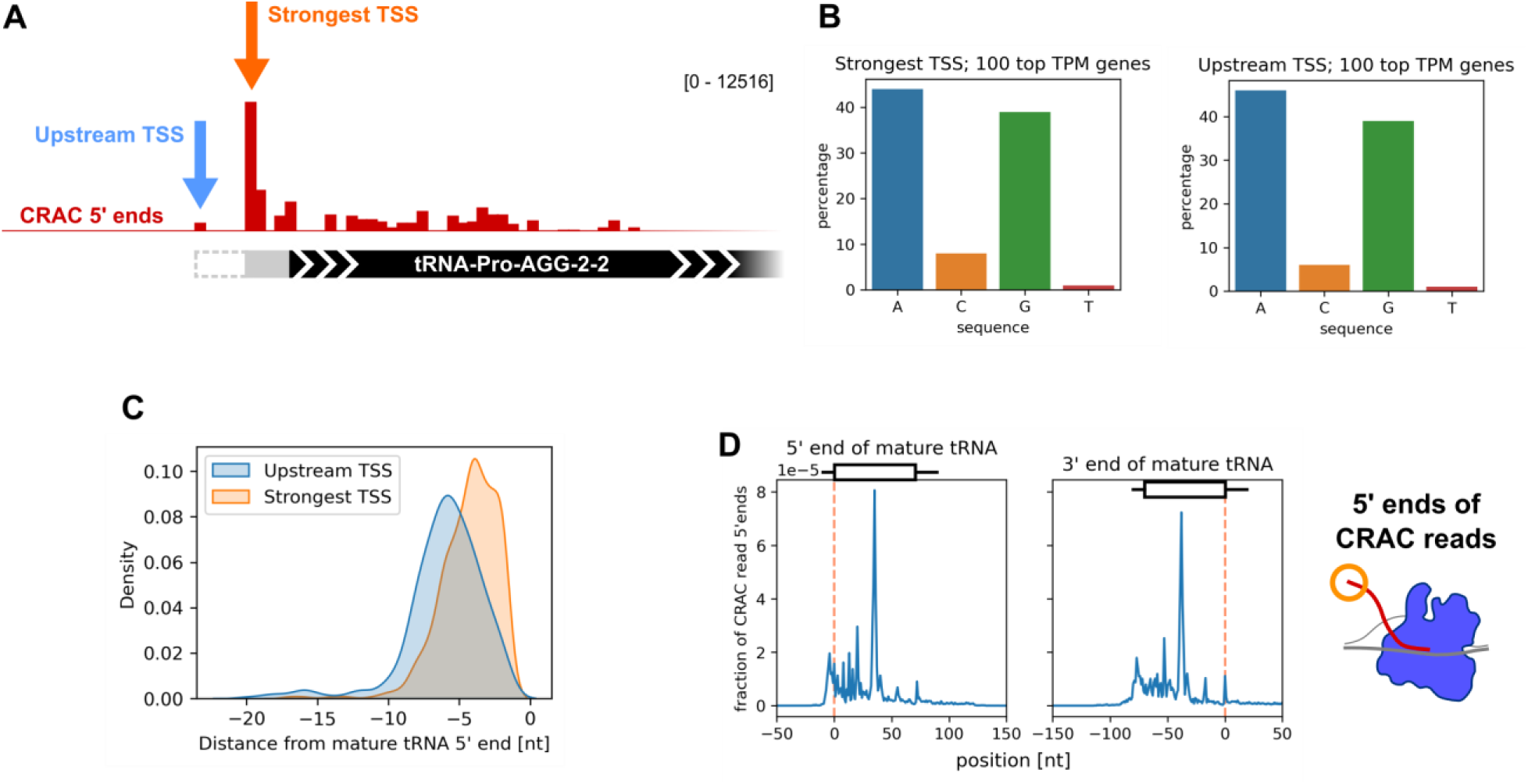
CRAC 5 ‘ends analysis reveals RNAPIII initiation is imprecise and tDNAs show signs of co-transcriptional processing. A: An example of upstream TSS in the tRNA-Pro-AGG-2-2 gene. TSS peaks were called by calculating local maxima of CRAC 5’ end coverage in the leader region. 5’ leaders resulting from initiation and the strongest and upstream TSS are indicated by a solid grey and dashed grey rectangle respectively. B: Nucleotide distributions of strongest and upstream TSSs. C: Kernel density estimate plots of TSS-mature tRNA 5’end distances for strongest and upstream TSSs. D: Metagene profiles of 5’ ends of CRAC reads on tDNAs aligned to the 5’ (left) or 3’ (right) end of the mature tRNA (position 0 on the x axis). Each line represents a biological replicate (n = 4).

At most tDNAs, the strongest TSS peak was preceded by a weaker signal located a few nucleotides upstream (Fig. 3C). Notably, we did not detect any TSS signal on the antisense strand, suggesting that RNAPIII initiation is strictly unidirectional. However, unlike RNAPI, which uses a single sharply defined TSS, RNAPIII initiation appears more flexible. We propose that this may reflect the architecture of the initiation complex, in which TFIIIC—composed of τA and τB subcomplexes connected by a flexible linker—first binds the B-box via τB. The τA subcomplex is then positioned at the A-box through a “fly-casting” mechanism^28^ and subsequently positions TFIIIB. Variable placement of τA, but also TFIIIB, might thus contribute to TSS heterogeneity.

We next asked whether CRAC data could reveal evidence of co-transcriptional tRNA processing, similar to cleavage or polyadenylation observed in other RNA polymerase systems. We searched for accumulation of 5′ ends at known processing sites. While we found no conclusive evidence for 5′ leader cleavage by RNase P (Fig. 3D, *left panel*), we detected a pronounced signal at the 3′ processing site recognized by RNase Z (Fig. 3D, *right panel*), suggesting that at least some 3′ end maturation may occur co-transcriptionally.

Together, these results reveal that RNAPIII initiation is unidirectional but not sharply defined, and suggest that the pre-tRNA processing event—3′ end cleavage—may occur co-transcriptionally. These findings add a new layer of complexity to tRNA biogenesis and underscore the importance of transcription initiation and nascent RNA processing in determining the sequence and structure of even short non-coding RNAs.

### The kinetics of RNAPIII are influenced by nascent RNA folding and thymidine stretches

Because the 3′ ends of CRAC reads correspond to the position of the RNAPIII active site, they provide a proxy for transcription kinetics: higher read density indicates slower elongation or pausing, whereas lower signal is associated with faster progression. Unexpectedly, RNAPIII distribution across tDNAs is non-uniform, with prominent coverage peaks accumulating along the gene body, suggesting variable elongation rates (Fig. 4A). While analysing profiles for intron-containing tDNAs (n = 34), we observed a marked drop in signal at the exon–intron boundary (Fig. 4B, top). We hypothesized that this sudden change in elongation rate may be caused by the folding of nascent RNA reflecting a T loop in mature tRNA, behind the polymerase, as previously proposed for RNAPI.^15^

**Figure 4.**
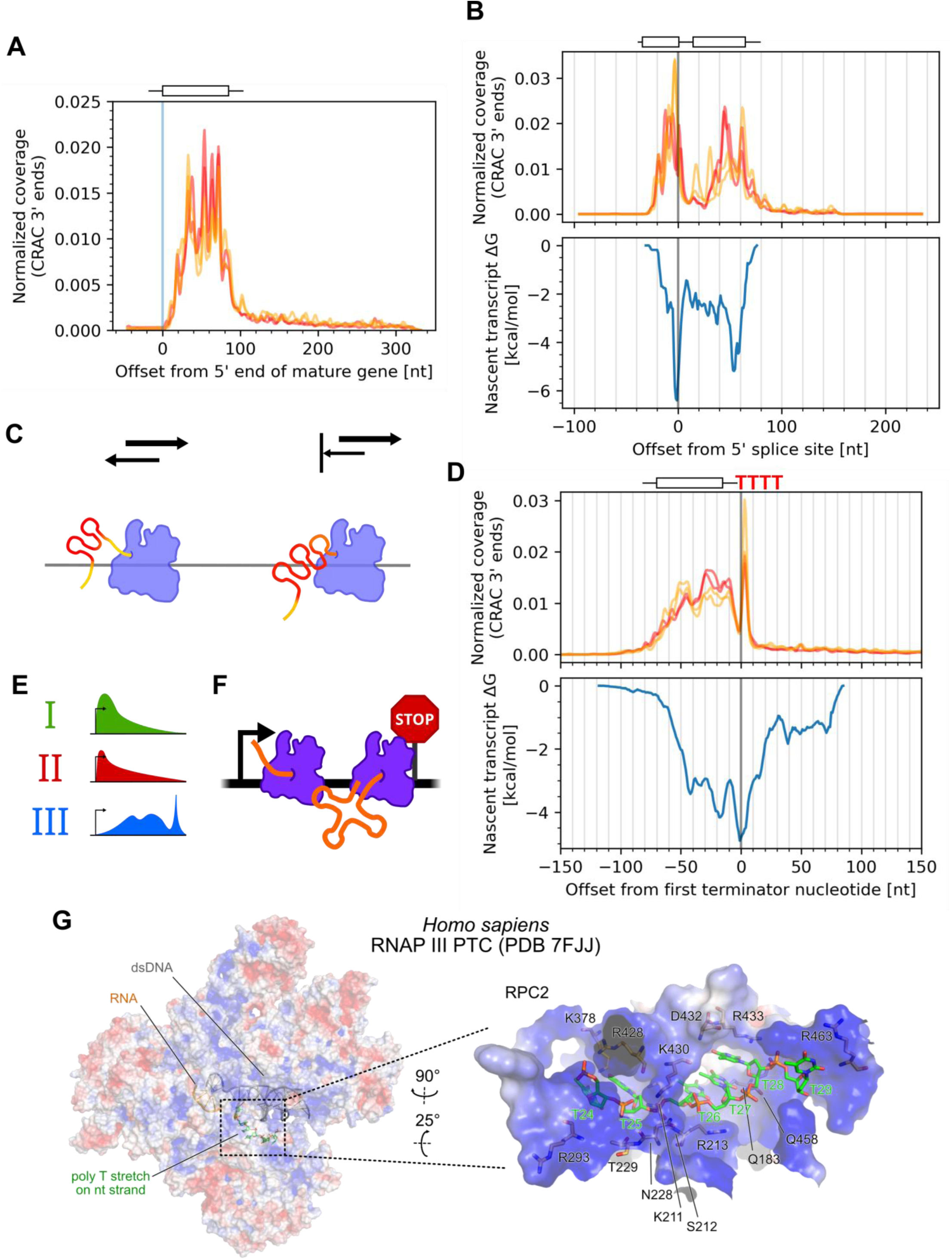
CRAC reveals RNAPIII transcription kinetics on tDNAs and pausing at thymidine stretches. A: Metagene profile of intron-free tDNAs aligned to the 5’ end of the mature tRNA (position 0 on the x axis). Each line represents a biological replicate (n = 4). B: Top: metagene profile of intron-containing tDNAs aligned to the last nucleotide of their 1^st^ exons (position 0 on the x axis). Each line represents a biological replicate (n = 4). Bottom: plot of nascent transcript free energy calculated in a 20 nt sliding window. C: Strong RNA secondary structures immediately behind the RNAPIII exit channel block backtracking and consequently increase elongation rate. D: Top: metagene profile of intron-free tDNAs aligned to the first terminator nucleotide (position 0 on the x axis). Each line represents a biological replicate (n = 4). Bottom: plot of nascent transcript free energy calculated in a 20 nt sliding window. E: Schematic representation of metagene polymerase density profiles for the three human RNA polymerases. F: Schematic representation of RNAPIII congestion on tDNAs due to termination-related RNAPIII pausing. G: Cryo-EM structure (PDB 7FJJ^29^) of human RNAPIII showing the terminator T-stretch binding to a positively charged patch on the polymerase complex.

To test this, we computed the folding free energy of the nascent transcript using a 20-nt sliding window, offset by 15 nt to account for the RNAPIII exit channel length (Fig. 4B, bottom). The resulting profile revealed that regions with more stable RNA structures (i.e., lower free energy) behind the RNAPIII correlate with drops in CRAC signal, indicating that nascent RNA folding promotes RNAPIII elongation. Similar correlation was observed for other, non-tRNA, RNAPIII transcripts (Fig. S5A). Given that RNAPs translocate via Brownian motion, we propose that RNA secondary structures forming behind the enzyme serve as barriers to backtracking, thus promoting the polymerase translocation forward (Fig. 4C), consistent with our previous work in yeast.^15^

We next focused on the 3′ ends of tDNAs, assuming that the prominent peak observed in this region corresponds to terminating RNAPIII. To test this, we aligned all intron-free tDNAs (n = 583) at the first nucleotide of their terminators. The resulting metagene profile again revealed variable elongation rates that correlated well with predicted free energy of nascent RNA folding (Fig. 4D)— with one notable exception at the termination site. Unique elongation dynamics at transcription termini have also been described for RNAPI and were attributed to co-transcriptional processing.^17^ In human RNAPIII, the sharp peak at the terminator suggests extremely slow elongation or pausing. This is consistent with previous findings in yeast, where RNAPIII is thought to enter a pre-termination complex upon encountering the terminator.^30^ Supporting this, structural studies of human RNAPIII revealed a positively charged tunnel through which the non-template DNA strand is funnelled, potentially aiding recognition of (T)-rich sequences (Fig. 4G and S5B).^31^ We also asked whether T stretches located elsewhere in the gene body—such as those present in the anticodon loops of lysine tRNAs—could similarly affect RNAPIII elongation (Fig. S5C). Indeed, the observed local deceleration confirms that T-stretches modulate RNAPIII kinetics even outside termination contexts, supporting the notion that recognition and interaction with such sequences by the polymerase is a general mechanism of elongation control.

Overall, the RNAPIII transcription profile appears distinct from those of RNAPI and RNAPII, with the most prominent peak consistently located at the terminator (Fig. 4E). This suggests that transcription termination may represent the most time-consuming step in the RNAPIII transcription cycle. Given the footprint of RNAPIII, a typical tDNA transcription unit can accommodate only two polymerase complexes between the TSS and the terminator (Fig. 4F). This spatial limitation supports the idea that slow termination could serve as a rate-limiting step in RNAPIII-mediated transcription.

### Purified RNAPIII recapitulates major features of *in vivo* elongation kinetics

To better understand and validate the kinetic features observed in the CRAC data, we performed *in vitro* elongation assays using purified human RNAPIII. We designed synthetic transcription scaffolds containing modular sequences, including structural elements (e.g., T-loops derived from the 3’ end of tDNAs), trailer and two terminators separated by a 6-nt spacer (Fig. 5A and S6A). In these assays, RNAPIII exhibited a pronounced tendency to pause at the final nucleotide of the terminator region (Fig. 5B–C). This observation was consistent with CRAC data, where RNAPIII accumulation was observed to shift progressively downstream with increasing terminator length (Fig. 5D and S6C).

**Figure 5.**
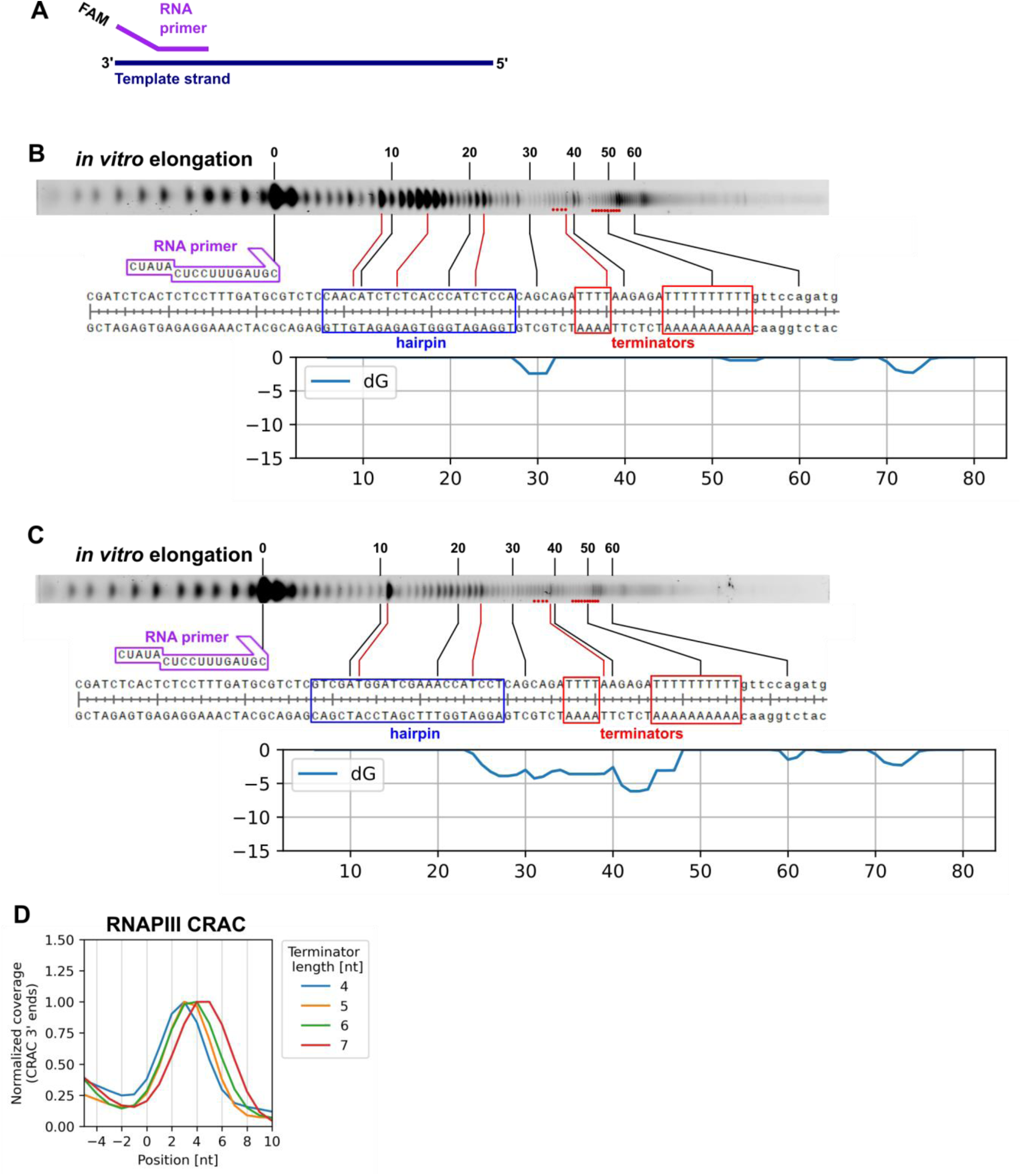
RNAPIII pauses on terminators *in vitro*. A: Schematic depiction of elongation assay scaffolds. B-C: RNAPIII elongation assays. Each panel presents a representative urea-polyacrylamide gel lane, the scaffold sequence, and free energy plot. The plots are aligned to the sequence. Blue boxes indicate the hairpin elements. Red boxes on the sequence and red dots on the gel images indicate terminator nucleotides. D: Terminator peaks from CRAC metagene profiles of tDNAs grouped by the length of their first terminator. Position 0 denotes the first terminator nucleotide. Plots were normalized to the peak value.

Importantly, these results indicate that terminator-associated pausing is not solely dependent on recognition of T-stretches in the non-template strand, consistently with previous data.^20^ Since the templates used in these assays were single-stranded, any observed effects must be attributed to nascent RNA features, within or outside the transcription bubble, rather than interactions with the non-template DNA strand. Supporting this, inclusion of a structured RNA element in the nascent transcript significantly enhanced elongation efficiency, as evidenced by reduced accumulation of paused polymerase along the scaffold (Fig. 5B-C).

The interplay between the termination sequence and nascent RNA folding has been discussed as a major mechanism for RNAPIII termination in yeast.^20,32,33^ Based on this, we hypothesized that the distance between a stable nascent tRNA structure and the terminator element can fine-tune termination efficiency in human: folding of the nascent RNA may promote translocation toward the terminator, where RNAPIII then stalls at the final nucleotide. Consistent with this model, we observed that highly stable RNA structures (low ΔG) can abolish termination-associated pausing (Fig. S6B).

### Termination of RNA polymerase III transcription is the rate-limiting step in tRNA synthesis

Despite these observations, the role of terminator length in regulating termination efficiency remained unresolved. To address this, we designed a dual-reporter plasmid system containing a synthetic tDNA followed by terminators of variable length, and an eGFP gene for internal normalization (Fig. 6A, S6D-E for detailed construct and primer design). These constructs were electroporated into K562 cells, and total RNA was harvested after 24 hours. Pre-tRNA and GFP expression were then quantified by RT-qPCR. Based on prior assumptions, we expected that longer terminators would enhance termination efficiency and lead to higher pre-tRNA levels. Instead, we observed the opposite: pre-tRNA expression dropped to ∼50% of the original level with terminators longer than 4T (Fig. 6B). This effect was most pronounced for the 6T terminator and persisted for terminators up to 12T. These findings suggest that RNAPIII termination impacts transcriptional output, but not via a simple linear enhancement model—rather, longer terminators appear to impede expression.

**Figure 6.**
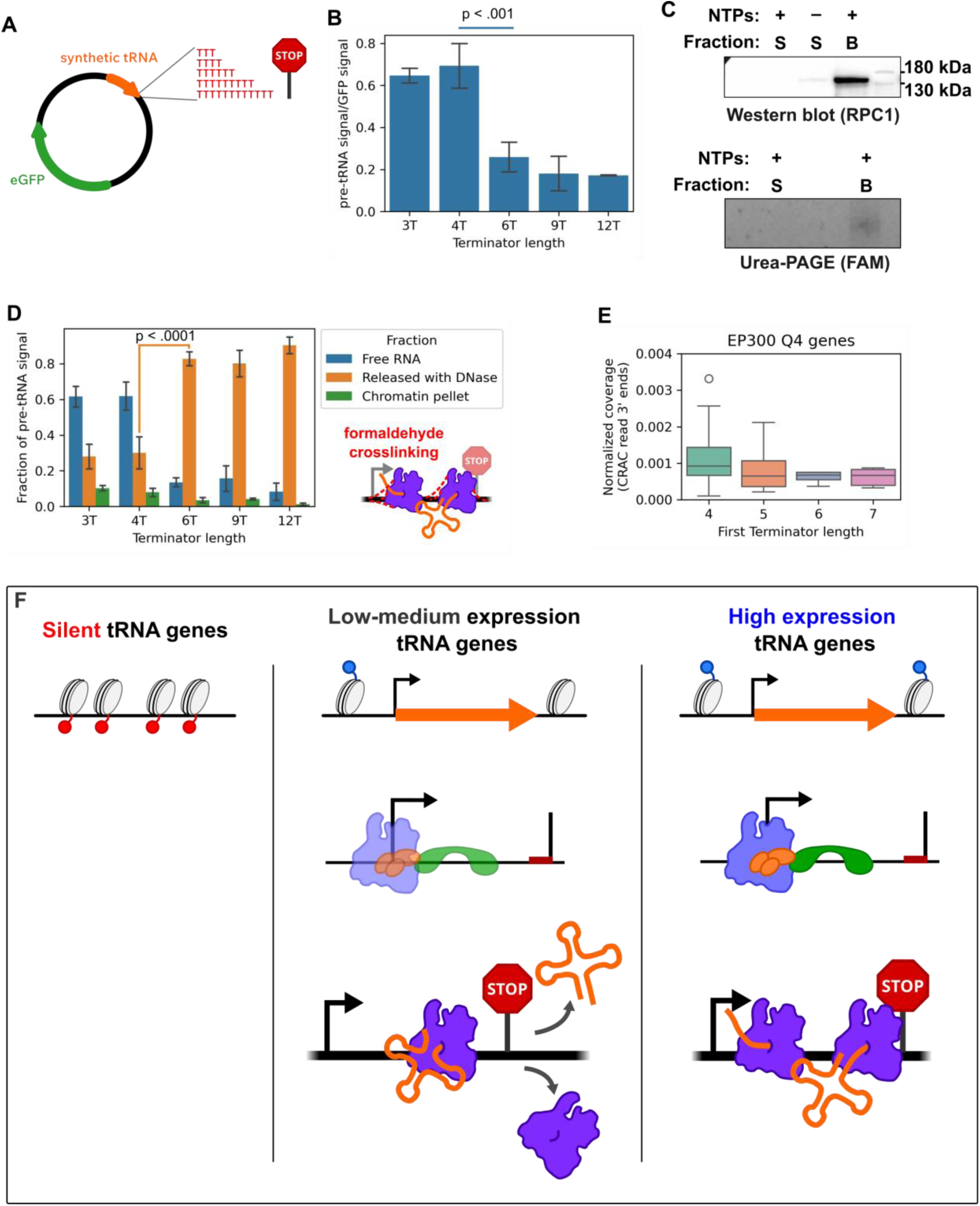
Terminator length influences transcript release by RNAPIII. A: Schematic depiction of the reporter system plasmids. B: tDNA expression from reporter plasmids measured by RT-qPCR. C: RNAPIII fails to release transcripts *in vitro*. Representative images of western blots and Urea-PAGE gels of bead and supernatant fractions after *in vitro* elongation assays are shown. S: supernatant fraction; B: bead-bound fraction. D: tDNA expression from reporter plasmids measured by RT-qPCR after formaldehyde crosslinking. E: tDNA expression measured by CRAC for highly expressed tDNAs, defined as those in the highest quartile of EP300 binding. F: Proposed model of tDNA expression regulation. Transcription is unable to initiate on silent genes because of closed chromatin (left). On tDNAs with low and moderate expression levels the chromatin is open and initiation factors TFIIIC and TFIIIB can bind to tDNA and recruit RNAPIII, which is free to elongate. Because initiation frequency is not saturated, there is no pressure for efficient transcript release (center). On highly expressed genes, chromatin is open due to favorable epigenetic modifications and/or factor binding. Initiation is efficient, saturating the tDNA with elongating RNAPIII. Because of short gene length, RNAPIII pausing at the terminator without releasing the transcript now presents a significant obstruction to efficient transcription. In this high-expression regime, termination becomes rate-limiting. p-values obtained using Student’s two-tailed t-test.

To assess transcript release, we adapted the *in vitro* elongation assay by using biotinylated DNA templates immobilized on streptavidin beads (Fig. S7A). After transcription, bead-bound and supernatant fractions were separated and analysed. Surprisingly, RNAPIII failed to release transcripts into the supernatant (Fig. 6C and S7B). Western blotting against RPC1 confirmed that the polymerase remained bound to the DNA template, suggesting that although RNAPIII paused at the terminator, it did not efficiently terminate or release the RNA in the assay.

This led us to propose that terminators trap RNAPIII *in vivo*, and longer terminators cause polymerase congestion on short transcription units and reduce expression. To test this, we repeated the reporter experiment, incorporating a formaldehyde crosslinking step prior to cell harvest to preserve RNAPIII–DNA and RNAPIII–RNA interactions on the reporter gene (Fig. 6E). Following chromatin pelleting, we separated the released RNA (Free RNA, blue). The chromatin pellet was then treated with DNase to release DNA-associated RNA (Released with DNase, orange), while the remaining pellet fraction was retained as chromatin pellet (green). RT-qPCR analysis showed that with a 4T terminator, most pre-tRNA signal was found in the free fraction. As terminator length increased, the free fraction signal decreased, while the DNA-associated fraction increased accordingly (Fig. 6D), confirming that longer terminators impair transcript release *in vivo*.

To validate these findings on endogenous genes, we turned again to our CRAC data. Genome-wide, tDNA expression levels do not clearly correlate with terminator length (Fig. S7C). However, when we focused on highly expressed tDNAs located in permissive chromatin regions (top quartile of EP300 ChIP-seq signal), we observed a trend consistent with our reporter assay: longer terminators were associated with lower expression (Fig. 6E). Finally, we calculated the thermodynamic stability of transcription bubbles at the terminator, by comparing the free energy of the RNA:DNA hybrid versus the reannealed DNA duplex (Fig. S7D). This analysis further supported a model of transcript release representing a rate-limiting step in RNAPIII transcription.

Taken together, these results—along with CRAC profiles and *in vitro* assays—demonstrate that during high-output tDNA transcription, RNAPIII termination is the rate-limiting step, a phenomenon not previously reported for any transcription system. Impaired or delayed transcript release leads to persistent polymerase pausing which, due to the short length of tDNAs obstructs efficient transcription initiation or re-initiation.

## DISCUSSION

In this study, we address a critical gap in the understanding of human transcription by providing the first transcriptome-wide evidence of RNAPIII engagement with tDNA. Until now, RNAPIII occupancy, typically measured by ChIP-seq, served as the best proxy for its transcription activity.^10,34^ Here, we applied the CRAC methodology, which enables direct detection of nascent transcripts associated with the RNAPIII complex. A key advantage of UV crosslinking in CRAC is its high signal-to-noise ratio, resulting from the covalent linkage between RNA and a tagged protein. This allows for stringent, denaturing washes and has previously been used to map protein interactions across all RNA classes ^11,15,35–37^. More recently, a similar approach was used to challenge earlier reports of *in vivo* RNA binding by chromatin proteins such as PRC2 and CTCF, demonstrating that such interactions are not detectable upon stringent purification protocol.^38^ Although UV crosslinking is less efficient (∼1%) than chemical fixation, it preferentially captures transient interactions. As a result, CRAC datasets are typically not as deep than those generated by other methods, but they provide uniquely informative data for studying transcription kinetics.

### Expression of RNPIII transcripts remains under multilevel control

In contrast to RNAPII transcripts, which are tightly regulated through a wide array of well-characterized mechanisms, relatively little is known about how RNAPIII-transcribed genes are controlled. For many years, tDNA expression was considered largely stable, with general repression mediated by the conserved transcription regulator Maf1.^39,40^ It was commonly assumed that the stoichiometry between different tRNAs is determined primarily by the number of gene copies encoding a given anticodon.^41^ However, more recent studies have identified a subset of “housekeeping tDNAs” that appear to remain constitutively expressed, suggesting a more nuanced regulatory landscape^9,12^.

In this study, we leveraged the K562 cell line—a well-characterized model from the ENCODE project^24^—to explore the regulation of RNAPIII activity in greater detail. Since mammalian tDNAs are frequently organized into genomic clusters, we examined whether this clustering is associated with coordinated expression of tDNAs within a locus. We found that most clusters contain genes of varying expression levels. While previous studies have implicated higher-order chromatin structure in the regulation of RNAPIII-transcribed genes^7,42^, our integrative analysis of RNAPIII CRAC data with high-resolution chromatin conformation (Hi-C) data^27^ yielded inconclusive results, suggesting that large-scale chromatin architecture may play a limited role in this aspect of RNAPIII regulation.

Our most informative findings emerged from comparative analyses of RNAPIII CRAC signals with chromatin modifications and transcription factor occupancy. As expected, RNAPIII activity correlated strongly with active chromatin marks such as H3K27ac and H3K4me2, consistent with previous observations^43,44^. Notably, RNAPIII occupancy as measured by ChIP-seq corresponded only partially with transcriptional activity detected by CRAC. Several features—SETDB1, c-Jun and ZUS12—consistently distinguished transcriptionally active RNAPIII loci, revealing previously unrecognized aspects of RNAPIII regulation. These findings suggest that RNAPIII activity is subject to complex, multilevel control beyond simple promoter binding or chromatin accessibility.

### RNAPIII presents dynamics opposite to the other RNA polymerases

RNA transcription proceeds through three main stages—initiation, elongation, and termination— each governed by distinct thermodynamic principles and regulatory mechanisms. For RNAPI and RNAPII, elongation is typically slowest near the 5′ ends of transcription units, as shown in multiple high-resolution studies.^15,45,46^ In contrast, our results reveal that human RNAPIII exhibits an opposing kinetic profile, with the slowest elongation occurring near the 3′ ends of its transcription units (Fig. 4E).

Transcription elongation is a dynamic process characterized by alternating regions of rapid and slow progression.^11,15,37,45,46^ All RNA polymerases share a basic mechanism for elongation that relies on a Brownian ratchet mechanism: nucleotide addition drives elongation without requiring external energy.^47^ The irreversible release of pyrophosphate during each nucleotide addition acts as a ratchet, ensuring directional movement and promoting transcript elongation. Because this process depends on Brownian motion—which is fast but inherently random—elongation is prone to oscillations, including both forward translocation and backtracking. Directionality is maintained by high concentrations of nucleotide triphosphates. Unlike RNAPII, which requires specialized factors to recover from backtracking, both RNAPI and RNAPIII possess intrinsic endonucleolytic activity within their core complexes, suggesting that backtracking and cleavage are integral to their transcription cycles.^48^

Even minimal forces applied in the reverse direction can stall RNA polymerases, highlighting the critical role of DNA supercoiling in regulating RNAP kinetics.^49–51^ Although current DNA supercoiling mapping methods lack the resolution to directly capture this effect, the widespread presence of RNA-DNA hybrids at tDNAs^52^ supports the idea that underwound DNA may affect RNAPIII processivity. Conversely, transcription can be accelerated locally by the folding of nascent RNA, which restricts backward translocation.^15,53^ Our data reveal a strong correlation between RNAPIII kinetics and nascent RNA folding, suggesting that—even for these short transcription units—folding of the emerging RNA exerts a major influence on polymerase dynamics, both *in vitro* and *in vivo*.

For RNAPII, nucleosome occupancy presents a significant barrier to elongation, leading to transcriptional pausing in both *in vitro* and *in vivo* systems.^37,45,47^ In contrast, tDNAs are typically located in nucleosome-depleted regions, with nucleosomes flanking but not occupying the transcription unit itself.^54^ This supports the view that actively transcribed RNAPIII loci are maintained in a chromatin environment that favours efficient elongation.

RNAPIII termination has been one of the most intensively studied aspects of this transcription system. It features several distinctive properties, including a facilitated recycling mechanism whereby RNAPIII can rapidly reinitiate transcription to maintain a high output from a given locus.^55^ Termination is mediated by canonical poly(T)-tracts in the non-template strand. In yeast, frequent read-through occurs unless the terminator contains at least seven uridines, which are required for efficient termination of tRNA transcription.^12^ More recent studies have shown that termination efficiency also depends on the sequence of the non-template strand^30^ and on structural elements within the transcription complex.^31^ Notably, work by Xie and colleagues^20^ in yeast demonstrated that RNA secondary structures can enhance Pol III termination. Interestingly, stalling at the 3′ end of the T stretch appears to be a unique feature of human RNAPIII. Our findings further extend these observations by showing that uridine residues within the gene body also modulate RNAPIII behaviour.

Most notably, we demonstrate that a terminator consisting of four uridines (4U) is optimal for human RNAPIII termination *in vivo*. This conclusion is supported by CRAC data, *in vitro* biochemistry and a reporter assay in which tDNA transcription is monitored at single loci. Surprisingly, we found that efficient termination cannot be evaluated solely by measuring read-through; it also depends on the release of the nascent transcript. While the precise mechanism remains unknown, we propose several, non-mutually exclusive possibilities:

1. RNAPIII pausing at T-rich sequences in the non-template strand remains essential but insufficient for termination;
2. Because nascent RNA folding influences RNAPIII kinetics, the spatial distance between the folded tRNA structure and the termination signal may regulate pausing efficiency;
3. A 4T sequence in the non-template strand may slow the polymerase just enough to allow shallow backtracking, enabling endonucleolytic cleavage and transcript release. In contrast, longer poly(T)-tracts might excessively stall the complex, thereby preventing efficient backtracking and cleavage and ultimately reducing termination efficiency.

#### Limitations of the study

This study used cell lines, which do not fully represent the complexity of human organisms, so RNAPIII regulation may differ in tissues *in vivo*. Although the POLR3A-tagged cell line exhibited normal growth, protein tagging—required for the CRAC protocol—could affect RNAPIII complex function or assembly. During RNAPIII CRAC analysis, we observed that increased signal may reflect prolonged transcript release rather than higher transcription rates; thus, CRAC readouts should not be considered direct measures of transcription rates. Still, we consider CRAC the best available indicator of gene transcriptional activity *in vivo*.

## Supplemental information

**Document S1.** Figures S1–S7

**Table S1.** Oligonucleotides used for RNAPIII tagging, reporter plasmid construction, in vitro elongation assays, and RT-qPCR assays

**Table S2.** Novel genes transcribed by RNAPIII

**Table S3.** Publicly available ENCODE ChiP datasets used in this study

## Data and code availability

- All CRAC sequencing data have been deposited at GEO: GSExxxxx and are publicly available as of the date of publication.
- Original RNA gel and western blot images have been deposited at Mendeley at DOI: 10.17632/ndn3mwt25f.1 and are publicly available as of the date of publication.
- Any additional information required to reanalyze the data reported in this paper is available upon request.

## Author Contributions

TWT and JM conceived the project and wrote the manuscript. JM, JW and TWT performed experiments. JM, AK, TWT and CM analyzed the data. All authors edited and reviewed the manuscript.

## Acknowledgements

We thank Magdalena Boguta, Christoph Engel and Daniel Schindler for critical reading of the MS and all members of Turowski Lab for stimulating discussion. We thank Prof. Daniel Schindler for sharing plasmids and generous guidance in establishing the MoClo system. TWT was supported by the Polish National Agency for Academic Exchange (PPN/PPO/2020/2/00004/U/00001), TWT, JM and AK were supported by National Science Center (2020/39/D/NZ2/02115). JM was supported by EMBO (SEG #10959) and Institute of Biochemistry and Biophysics Intramural grant FBW-SD-03/2024.

## Methods

### EXPERIMENTAL MODEL and SUBJECT DETAILS

#### Cell lines

K562 cell line expressing C-terminal 8xHis-FLAG tagged RPC1 subunit was created using CRISPR-Cas9 genome editing utilizing homology-directed repair. Guide RNA and repair template sequences are listed in Table S1. Successful tagging was confirmed with PCR, Sanger sequencing and immunoprecipitation experiments.

### METHODS DETAILS

#### *In-vivo* RNA crosslinking

K562 POLR3A-HF cells were grown in RPMI-1640 medium supplemented with 10% FBS and Antibiotic/Antimycotic. Upon reaching culture density of ∼0.8·10^6^ cells/ml, cell suspensions were centrifuged, resuspended in PBS and cross-linked using the VariX crosslinker at 400 mJ·cm⁻² and wavelength of 254 nm. Cells were centrifuged and the pellets were snap frozen in liquid nitrogen and stored at −80°C. Approximately 0.8·10^9^ cells were used per sample.

#### CRAC

Samples were processed as previously described.^11^ However, phosphatase treatment was omitted, so the 3’-OH ends required for linker ligation are present only on nascent RNA transcripts. Cells were lysed by resuspending in lysis buffer (50 mM Tris-HCl pH 7.5, 0.1 M NaCl, 1% IGEPAL CA-630, 5 mM MgCl_2_, 0.5% sodium deoxycholate, 0.1% SDS) and incubating on ice for 10 minutes. 20U of RQ1 DNase per sample was added and the samples were incubated at room temperature for 10 minutes, then centrifuged at 2100 g for 5 minutes at 4°C. The cleared lysates were incubated with M2 antiFLAG magnetic beads for two hours at 4°C, with nutation. Subsequently, the beads were washed three times with lysis buffer, then two times with buffer C (50 mM Tris-HCl pH 7.8, 50 mM NaCl, 0.1% IGEPAL CA-630). Protein:RNA complexes were treated with RNace-It for 10 minutes at 23°C with shaking. The beads were washed once with lysis buffer, then FA2 buffer (50 mM HEPES/KOH pH 7.6, 500 mM NaCl, 1 mM EDTA, 1% Triton X-100, 0.1% sodium deoxycholate) and FA3 buffer (10 mM Tris/HCl pH 7.8, 250 mM LiCl_2_, 1 mM EDTA, 0.5% NP40 / IGEPAL, 0.5% sodium deoxycholate), then twice with buffer C. Complexes were eluted by incubating two times with 3xFLAG peptide in lysis buffer for 5 minutes at 37°C with shaking. The 400 μl eluate was adjusted for nickel affinity purification with the addition of 400 mg guanidine hydrochloride, 45μl NaCl (3M) and 3μl imidazole (2.5M) and added to 75 μl of washed nickel beads. The nickel beads were incubated at room temperature for 1 hour, then at 4°C for 1 hour. Following incubation, the nickel beads were washed three times with buffer B1 (6.0M guanidine hydrochloride, 50mM Tris-HCl pH7.5, 500mM NaCl, 0.1% Triton-X100, 10mM imidazole), and two times with C buffer. Beads were then transferred to spin columns. Subsequent reactions (80μl total volume for each) were performed in the columns, and afterward washed once with B1 and three times with buffer C: 1. 3’ linker ligation (1x PNK buffer (NEB), 10% PEG8000, 20U T4 RNA Ligase II truncated K227Q, 80U RNasIn, 80pmol pre-adenylated 3’ linker 16°C overnight). 2. 5’ end phosphorylation and radiolabeling (1x PNK buffer (NEB), 40U T4 PNK (NEB), 40μCi 32P-γATP; 37°C for 45 min, with addition of 100nmol of ATP after 30 min). 3. 5’ linker ligation (1x PNK buffer (NEB), 10% PEG8000, 40U T4 RNA ligase I (NEB), 80U RNasIN, linker, 200pmol 5’ linker, 1mM ATP; 16°C for 3h, then 25C for 2h). The beads were washed four times with buffer B1. Protein:RNA complexes were eluted in 2×50 μl of elution buffer (6.0M guanidine hydrochloride, 50mM Tris-HCl pH7.5, 500mM NaCl, 0.1% Triton-X100, 10mM imidazole, 300 mM imidazole) and ethanol precipitated for 1h at −80°C. RNPs were pelleted at 21,000g for 20min at 4°C, washed in cold acetone and resuspended in 1X NuPAGE sample loading buffer supplemented with 8% β-mercaptoethanol. The sample was denatured by incubation at 65°C for 10min, and run on a 4%–12% Bis-tris NuPAGE gel at 130V. The protein:RNA complexes were transferred to Hybond-C nitrocellulose membranes with NuPAGE MOPS transfer buffer with 10% methanol for 1.5h at 100V. Labelled RNA was detected by autoradiography. The appropriate region was excised from the membrane and treated with 0.2μg/μl Proteinase K (50mM Tris-HCl pH7.5, 50mM NaCl, 0.5%SDS, 1mM EDTA; 2hr 55°C with shaking) in a 400μl reaction. The RNA component was isolated with a standard phenol:chloroform extraction followed by ethanol precipitation with 1μl of GlycoBlue. The RNA was reverse transcribed using Superscript IV for 15 minutes at 50°C in a 20μl reaction. The reactions were then cooled to room temperature and transferred to ice for 3 minutes. 2 ul of ExoI nuclease per sample, and incubated for 30 minutes at 37°C, then 20 minutes at 80°C. The resulting cDNA was amplified by PCR in 50μl reactions using Phusion DNA polymerase (2 μl template, 22-25 cycles). PCR reactions were combined, precipitated in ethanol and resolved on a 3% Metaphore agarose gel. A region corresponding to 140 to 200 bp was excised from the gel and extracted using the Min-elute kit. Libraries were measured with Qubit and sequenced using Illumina NextSeq 2000 with 100bp single end reads.

#### Purification of human RNAPIII and in vitro elongation assays

Human RNAPIII was purified as described in ^31^. For elongation assays, 10 pmol each of template strand DNA and FAM-labeled RNA primer were incubated in hybridization buffer (20 mM HEPE pH 7.5, 6 mM MgCl_2_, 100 mM NaCl, 10 mM DTT) for 3 minutes at 95°C, then cooled down to 20°C at a rate of 0.1°C/s in a thermocycler. 2 pmol of resulting template-primer scaffold was mixed with 2 pmol purified RNAPIII and incubated at 20°C for 10 minutes. Transcription was then started by adding an equal volume of 2 mM NTPs in nucleotide buffer (20 mM HEPES pH 7.5, 60 mM (NH_4_)_2_SO_4_, 10 mM MgSO_4_, 10 mM DTT) for a final concentration of 1 mM of each NTP. Samples were incubated at 37°C for 10 minutes, after which the reaction was stopped by adding formamide-containing 2x RNA loading dye. Samples were then resolved on 12% Urea-polyacrylamide gels and visualized using a Typhoon laser scanner.

For reactions performed on streptavidin beads, the scaffold-RNAPIII complex was incubated with beads at 20°C for 30 minutes, after which the elongation assay was performed as described above. The bead and supernatant fractions were separated and analysed separately.

To assess RNAPIII falloff, samples from both bead and supernatant fractions were taken and analysed using western blotting. Bead samples were directly denatured and loaded on gel, while supernatant samples were TCA precipitated beforehand. After electrophoresis proteins were transferred from the gel onto a nitrocellulose membrane, then probed with an anti-RPC1 monoclonal antibody following the manufacturer’s protocol. The membrane was then visualized using a HRP-conjugated secondary antibody.

#### In vivo reporter system construction

The reporter plasmid was constructed using the MoClo system described in ^56,57^. A synthetic tRNA construct was created by first amplifying a genomic fragment containing the tRNA-Arg-TCG-4-1 tDNA from WT K562 cells. The resulting amplicon was used as a template for PCR with primers containing MoClo overhangs for subsequent assembly. The reverse primers were designed to also replace the 3’ trailer sequence with a complementary one to make the synthetic pre-tRNAs distinct from endogenous ones, as well as modify the oligo-(T) terminator length. These constructs were then one-step assembled with flanking sequences amplified from mouse genome, as well as eGFP and KanR genes cloned from the pMax_GFP plasmid.^58^

#### In vivo reporter assays

WT K562 cells growing in logarithmic phase were transfected with 1 μg of reporter plasmid using the Lonza Nucleofector II electroporator. 1·10^6^ cells were used per transfection. 24 hours post-transfection, cells were washed in PBS, harvested by centrifugation and total RNA was extracted using TRI reagent.

For experiments with chromatin fraction separation, 24 hours post-transfection, 1·10^6^ cells were resuspended in 10 ml growth medium and crosslinked by adding 270 μl of 37% formaldehyde and incubating on a rotator for 5 minutes at room temperature. Crosslinking was then quenched by adding 1 ml of 2.5M glycine and incubating for further 2 minutes. Afterwards cells were centrifuged, washed with PBS, pelleted, snap frozen in liquid nitrogen and stored at −80°C.

Cells were gently thawed and lysed in lysis buffer (50 mM Tris-HCl pH 7.5, 0.1 M NaCl, 1 % IGEPAL CA-630, 5 mM MgCl_2_, 0.5% sodium deoxycholate, 0.1% SDS) with cOmplete protease inhibitor cocktail by incubating on ice and gently pipetting with a wide-bore pipette tip. The lysates were then centrifuged for 3 minutes at 3000g at 4°C. The supernatant was removed and labelled as the *Free RNA* fraction. The chromatin pellet was then resuspended and washed three times in wash buffer (40 mM Tris-HCl pH 7.5, 10 mM MgCl_2_, 1 mM CaCl_2_), then resuspended in 50 μl 1X RQ1 DNase buffer containing 1 μl RQ1 DNase and 1 μl RNasin, then incubated at 37°C for 30 minutes to release chromatin-bound DNA. After centrifugation, the supernatant was removed and labelled as the *Released with DNase* fraction. The remaining pellet was resuspended in wash buffer and labelled as the *Chromatin pellet* fraction. RNA was then isolated from all fractions using GTC-phenol extraction.

For all samples, reverse transcription was performed with random hexamers, using 200-1000 ng of RNA after DNase treatment with RQ1 DNAse and SuperScript III reverse transcriptase, according to the manufacturer’s protocol. RT-qPCR assays were performed on a Roche LightCycler 480 instrument.

### QUANTIFICATION AND STATISTICAL ANALYSIS

#### Pre-processing and data alignment

Illumina sequencing data were demultiplexed using in-line barcodes and in this form were submitted to GEO. First quality control step was performed using FastQC software (http://www.bioinformatics.babraham.ac.uk/projects/fastqc/) considering specificity of CRAC data. Raw reads were collapsed to remove PCR duplicates using FASTX-collapser v0.0.14 (http://hannonlab.cshl.edu/fastx_toolkit/) then inline barcodes were removed using pyBarcodeFilter.py script from pyCRAC package v3.0.^59^ The 3’ adaptors were removed using flexbar v3.5.0 ^60^ with parameters -at 1 -ao 4 –u 3, and filtered to retain only reads containing the 3’ adaptor.

All datasets were aligned to the hg41 human genome using STAR v2.7.10a.The 3’ ends or the 5’ ends of reads were selected using an in-house awk script and 1 nt resolution BigWig files were generated using bamCoverage script from deepTools v3.5.1 package.^61^ Sam file operations were performed using SAMtools v1.15.1. ^62^

#### ChIP data processing and analysis

Raw sequencing reads were downloaded from GEO and processed using a custom Snakemake (v7.32.4) pipeline. Initial alignment and sorting were performed with Bowtie2 (v. 2.3.4.3) and Samtools (v.1.9), aligning reads to the reference genome hg41 and generating sorted BAM files. Duplicate reads were marked using Sambamba (v. 0.6.6). Marked duplicates were subsequently removed using Samtools.

Peak calling was conducted using MACS3 (v. 3.0.0a6) with the --bdg option to generate signal tracks. Input and IP samples were grouped by condition, and peaks were called using a genome size parameter (-g hs) appropriate for human data. The resulting .narrowPeak files were converted to BED4 format by extracting relevant columns (chromosome, start, end, and signal value) using python Pandas. These BED files were then transformed into BigWig format using bedGraphToBigWig, with chromosome sizes provided via the chrom parameter.

Subsequent steps were executed in Python Jupyter (v. 5.8.1) notebooks to ensure reproducibility and analysis transparency.

#### Hi-C data processing and analysis

Raw data were processed into .cool files using the Distiller Snakemake (v7.32.4) pipeline developed by Open2C (https://github.com/open2c/distiller-sm). Subsequent analyses were performed with Cooltools v0.7.1 ^63^, which was used to generate insulation boundaries from Hi-C data. These boundaries were then applied to examine tDNA expression within specialized clusters.

#### Linear tDNA cluster calling

A linear tDNA cluster was defined as a group of at least 2 tDNAs separated by an intergenic distance lower than a chosen cutoff value. A distance cutoff of 10 kbp was used for the final analysis and main text figures. Figures for smaller distances are shown in Figure S4A.

#### RNA polymerase III profiles

Downstream analyses were performed using Python 3.10 Jupyter notebooks, using existing libraries as well as in-house scripts and the trxtools v0.3.0 package (https://github.com/TurowskiLab/trxtools).

##### Folding of nascent RNA

Each sequence was divided into segments using a rolling window of w nt, where w was the length of RNA considered to form structure (chosen 20 nt, tested range 10-40 nt). The folding energy at 37°C was calculated using hybrid-ss-min from UNAfold package v3.8 (Markham and Zuker, 2008). Folding energies were associated with the position of last nucleotide in the sequence and a 15 nt offset was applied to exclude the 3’ end of the nascent RNA inside the RNAP exit channel and calculate folding energy only for the extruded RNA.

##### Statistical analyses and numerical methods

All plots and statistical analyses of this work were performed using Python 3.10 Jupyter notebooks and python library scipy v1.10.1.

