## Supplementary materials for "High-resolution mapping of human RNA polymerase III reveals transcription termination as a rate-limiting step"

### SUPPLEMENTARY MATERIAL

**Figure S1.** (related to Figure 1)
**Figure S2.** (related to Figure 2)
**Figure S3.** (related to Figure 2)
**Figure S4.** (related to Figure 2)
**Figure S5.** (related to Figure 4)
**Figure S6.** (related to Figure 5)
**Figure S7.** (related to Figures 5 and 6)

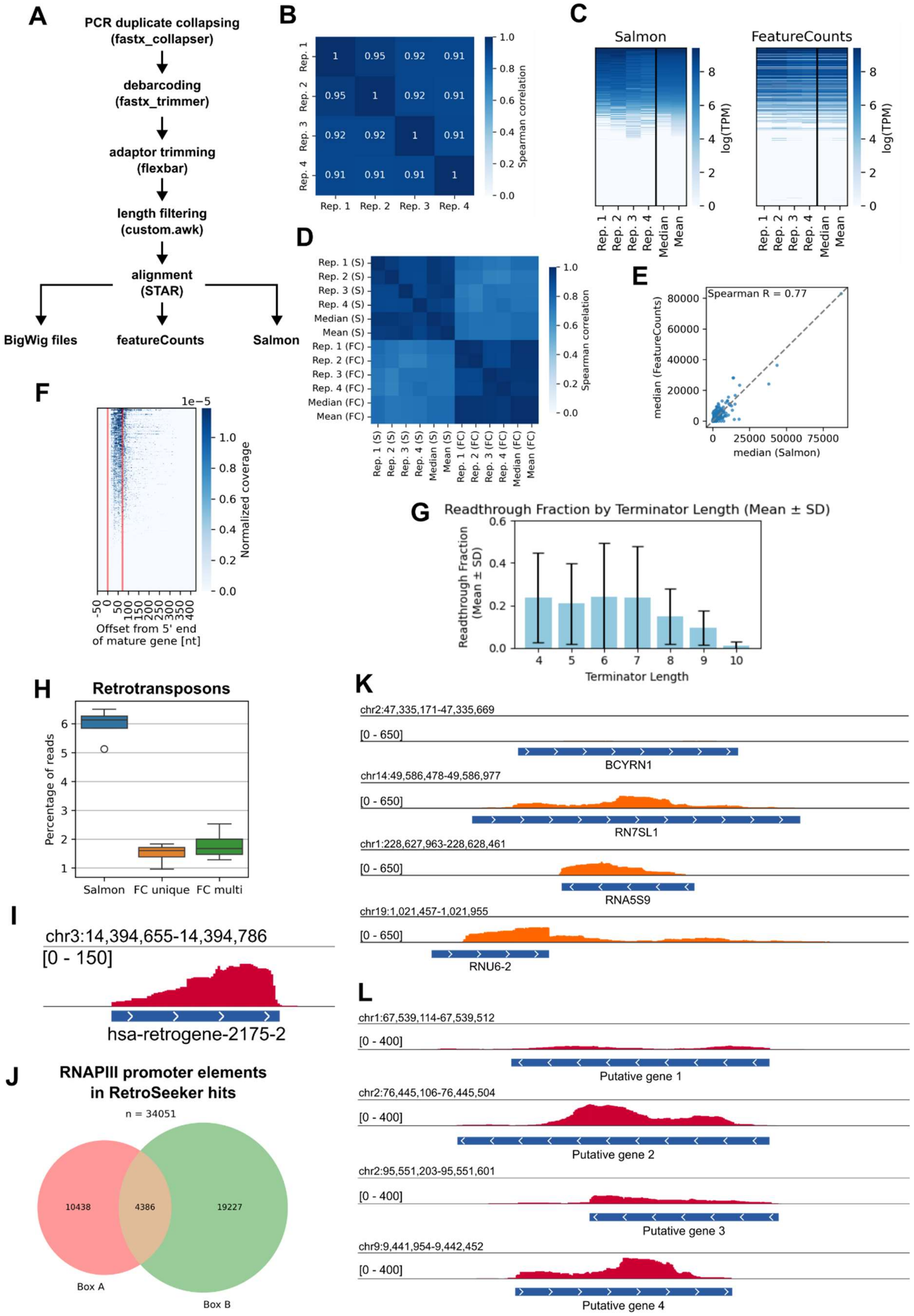


**Figure S1. Mapping CRAC reads to tDNAs, retrotransposons and novel genes, related to Fig 1**

A: Outline of CRAC data preprocessing steps.

B: Spearman correlation heatmap of RNAPIII CRAC libraries.

C: Heatmaps of tRNA expression measured by Salmon (mapping-based mode) and featureCounts.
D: Spearman correlation heatmap comparing tRNA expression measurements obtained with Salmon (S) and featureCounts (FC).
E: Scatterplot of expression values for every tRNA gene obtained with Salmon and featureCounts.
F: Heatmap showing profiles of CRAC read 3' end coverage across tDNAs.
G: Bar plot of readthrough fraction, calculated as the fraction of tDNA coverage past the primary terminator, in tDNAs with different primary terminator lengths.
H: Share of retrotransposon transcription in total RNAPIII transcription quantified with Salmon and featureCounts (FC) with multimapping disabled (*unique*) and allowed (*multi*).
I: Example gene browser track showing CRAC coverage of a retrotransposon element.
J: Counts of retrotransposons with detected RNAPIII type 2 promoter elements.
K: Non-tRNA RNAPIII transcripts detected using CRAC: BC200 RNA (BCYRN1), 7SL RNA (RN7SL1), 5S rRNA (RNA5S9), U6 snRNA (RNU6-2)
L: Examples of unannotated RNAPIII genes. Putative genes 1-3 reside in intergenic regions. Gene 4 is located in an intron of PTPRD gene in an antisense orientation.

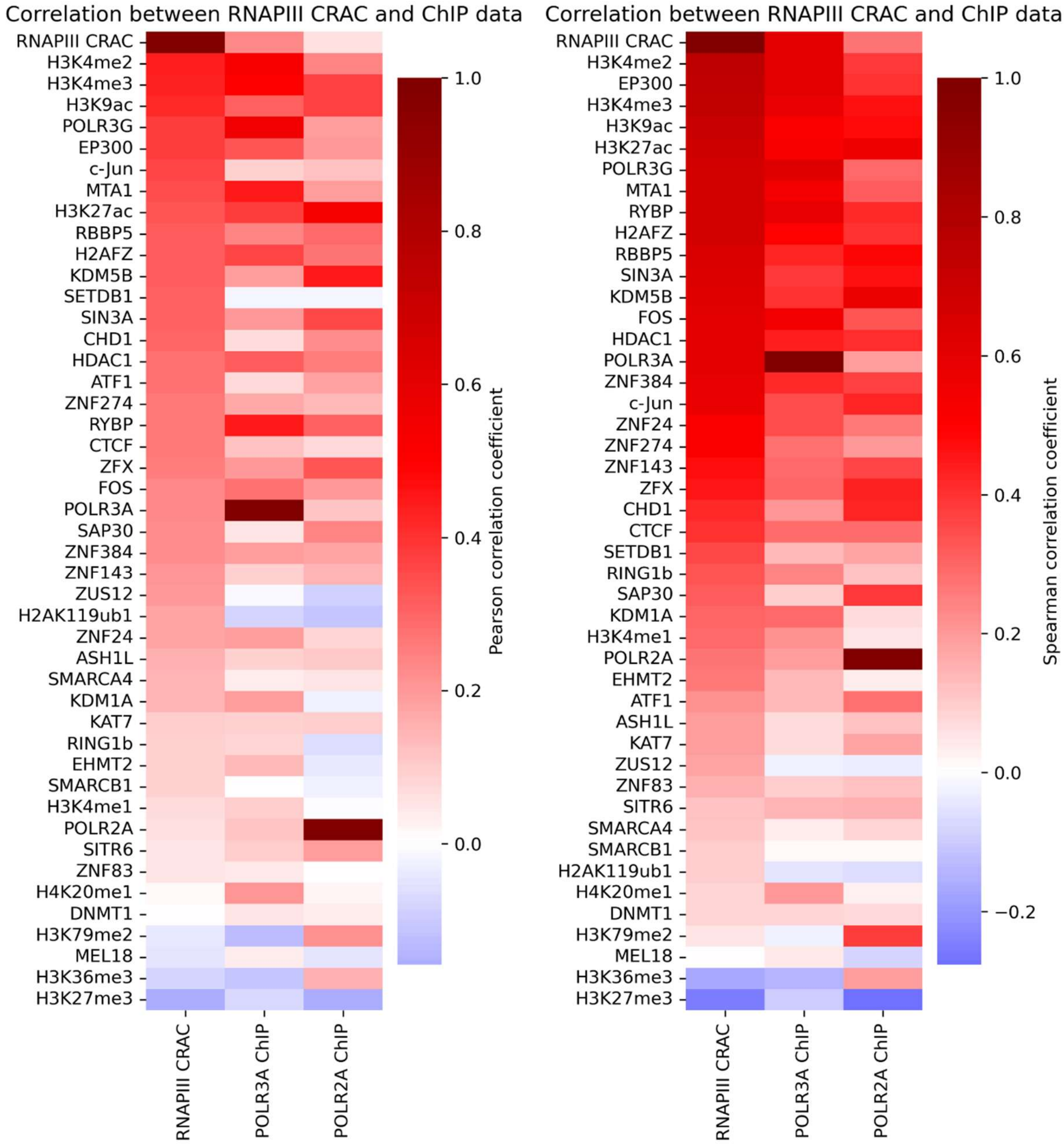


**Figure S2. Correlation between CRAC and ChIP datasets, related to Figure 2**

Differential heatmaps of chromatin-associated factor and mark binding. Pearson (left) and Spearman (right) correlation factors are shown.

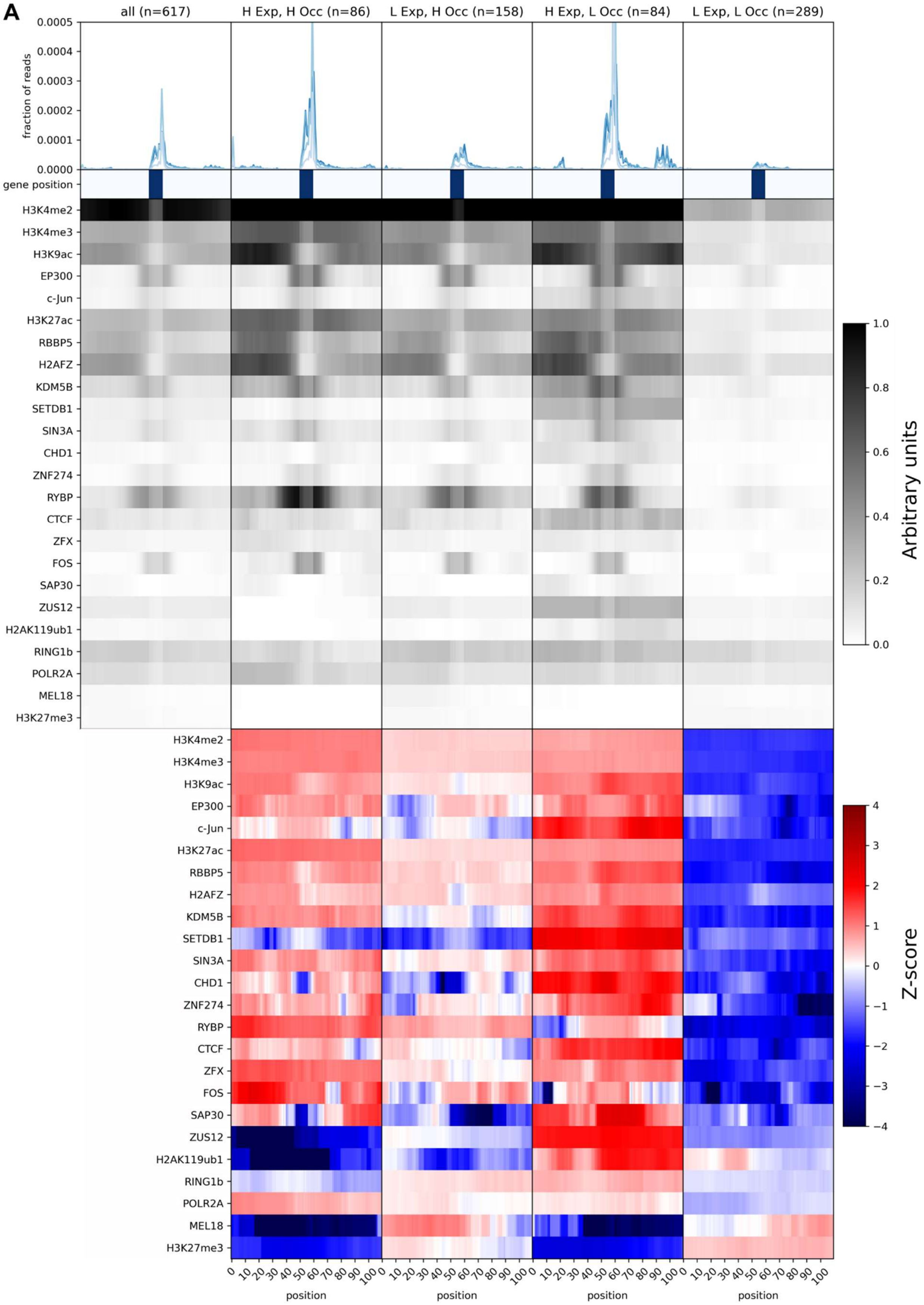
A
all (n=617)
H Exp, H Occ (n=86)
L Exp, H Occ (n=158)
H Exp, L Occ (n=84)
L Exp, L Occ (n=289)
fraction of reads
gene position
H3K4me2
H3K4me3
H3K9ac
EP300
c-Jun
H3K27ac
RBBP5
H2AFZ
KDM5B
SETDB1
SIN3A
CHD1
ZNF274
RYBP
CTCF
ZFX
FOS
SAP30
ZUS12
H2AK119ub1
RING1b
POLR2A
MEL18
H3K27me3
Arbitrary units
Z-score
position

**Figure S3. Chromatin-associated factors and epigenetic marks are high-level tDNA expression regulators, related to Fig 2**

A: Chromatin associated factor binding at tDNAs. tDNAs were grouped according to their expression and RNAPIII occupancy as in Fig 2B-C. Top panel: metagene profiles of CRAC read 5’end coverage. Middle panel: ChIP occupancy profiles. Bottom panel: heatmaps of occupancy differences between a given group and the mean for all tDNAs, expressed as Z-scores.

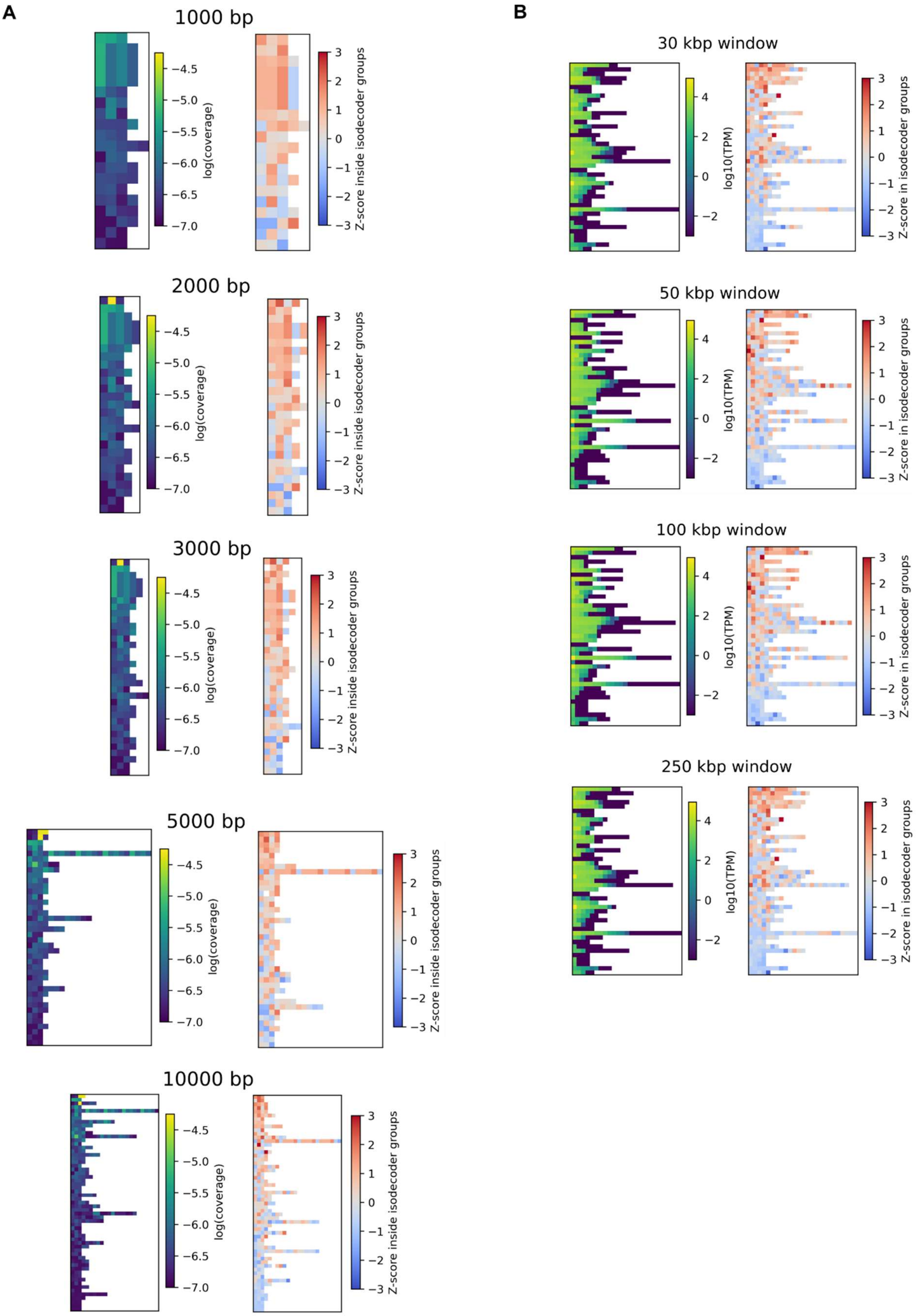


**Figure S4. tDNA expression in linear clusters and topological domains, related to Figure 2**
A: tDNA expression in linear clusters. Absolute expression (left heatmap in every pair) was measured as log(10) of CRAC read 3' end coverage. Expression of every gene relative to its isoacceptor family

was calculated as a Z-score in that group (right heatmap in every pair). Numbers above every heatmap pair indicate the maximum permitted distance between genes in a single cluster. Clusters containing less than three genes are not shown.
B: tDNA expression in topological domains. Expression was calculated as in A. Window size used to determine domain boundaries (see Methods section for details) is indicated above each heatmap pair.

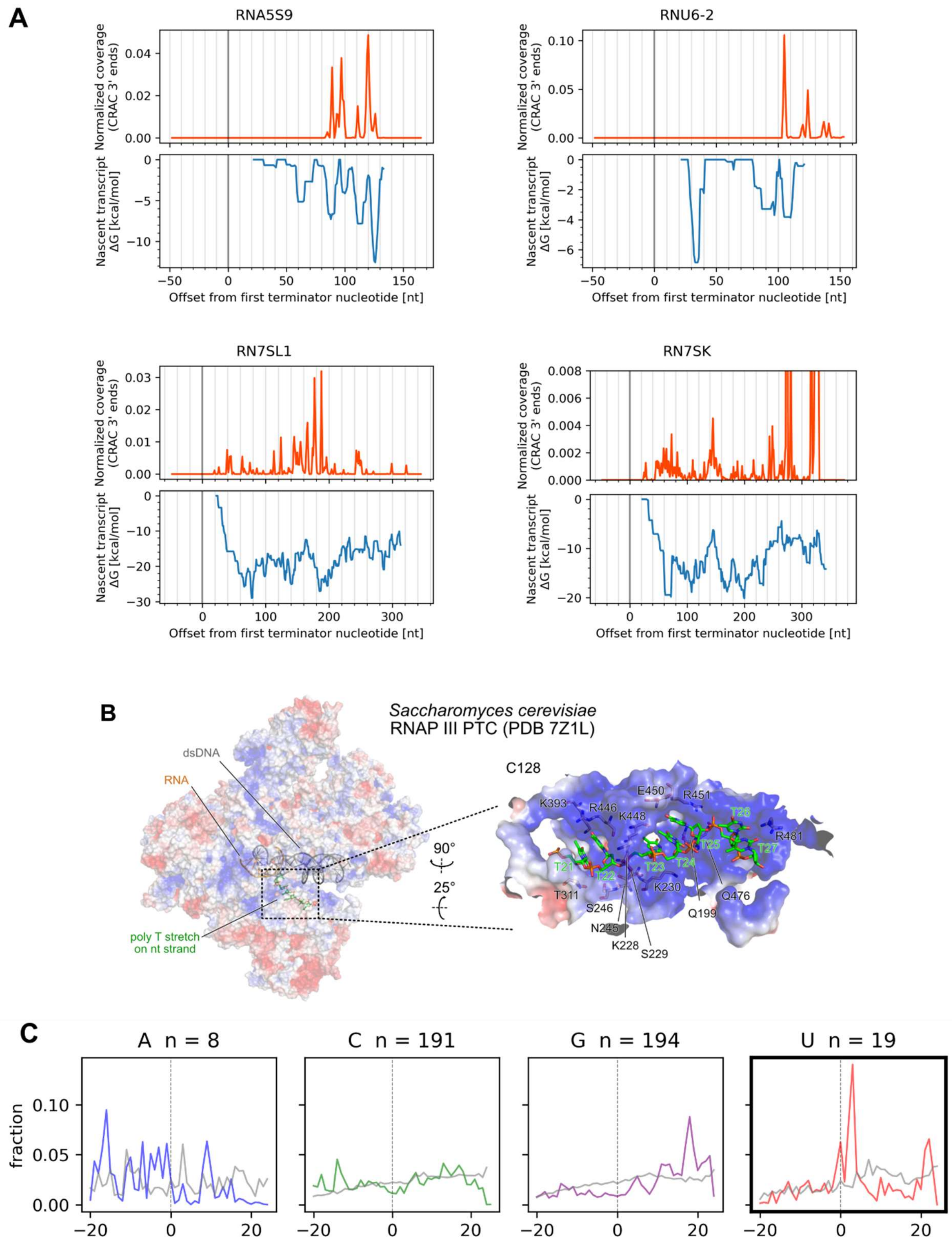


**Figure S5. RNAPIII transcription kinetics, related to Figure 4**

A: Top in every panel: Metagene profiles of 5S rRNA, U6 snRNA, 7SL RNA, and 7SK RNA genes aligned to the 5' end of the mature transcript (position 0 on the x axis). Each plot represents the median of 4 biological replicates. Bottom in every panel: plot of nascent transcript free energy calculated in a 20 nt (5S, U6) or 50 nt-wide (7SL, 7SK) sliding window.

B: Cryo-EM structure of *Saccharomyces cerevisiae* RNAPIII showing the terminator T-stretch binding to a positively charged patch.

C: Metagene profiles centered around nucleotide stretches. Number of genes containing a given nucleotide stretch is indicated above each panel. Colored lines: mean coverage between 4 biological replicates. Grey lines: coverage sampled from random regions in tRNAs containing indicated nucleotide stretches. Position 0 on the x axis indicates the first nucleotide of a stretch.

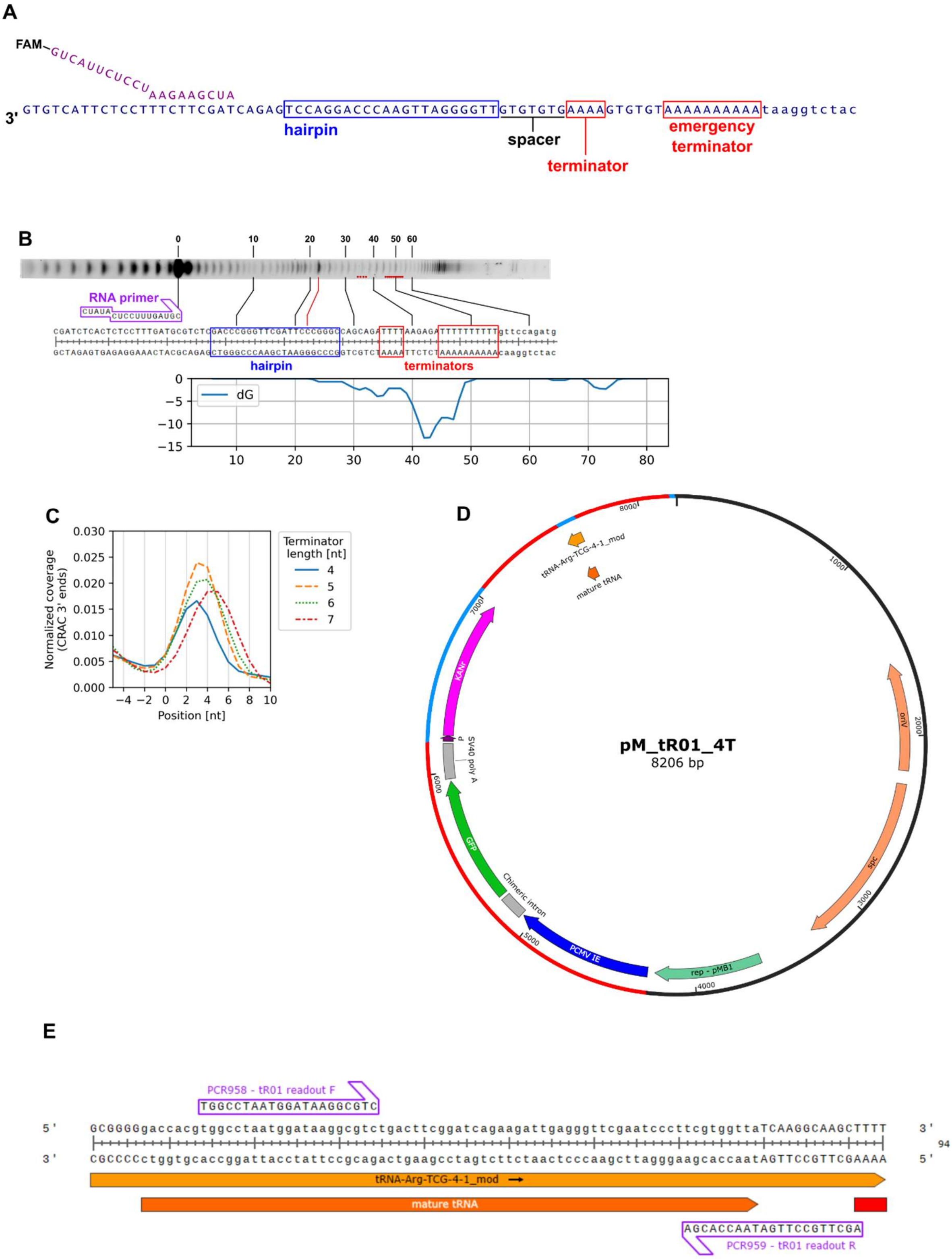


**Figure S6. In vitro and in vivo assays, related to Figure 5**

A: Example sequence of scaffolds used for *in vitro* elongation assays.

B: RNAPIII elongation assay with strong structural element. Shown are the urea-polyacrylamide gel lane, the scaffold sequence, and free energy plot aligned to the sequence. Blue box indicates the hairpin element. Red box on the sequence and red dots on the gel images indicate terminator nucleotides.

C: Terminator peaks from CRAC metagene profiles of tRNA genes grouped by the length of their first terminator. Position 0 denotes the first terminator nucleotide.
D: Example of plasmid used for in vivo reporter assays. See Methods section for details on plasmid construction.
E: Sequence of synthetic tDNA used for reporter assays, here depicted with a 4T terminator (red box below sequence). Annealing sites for readout primers are indicated.

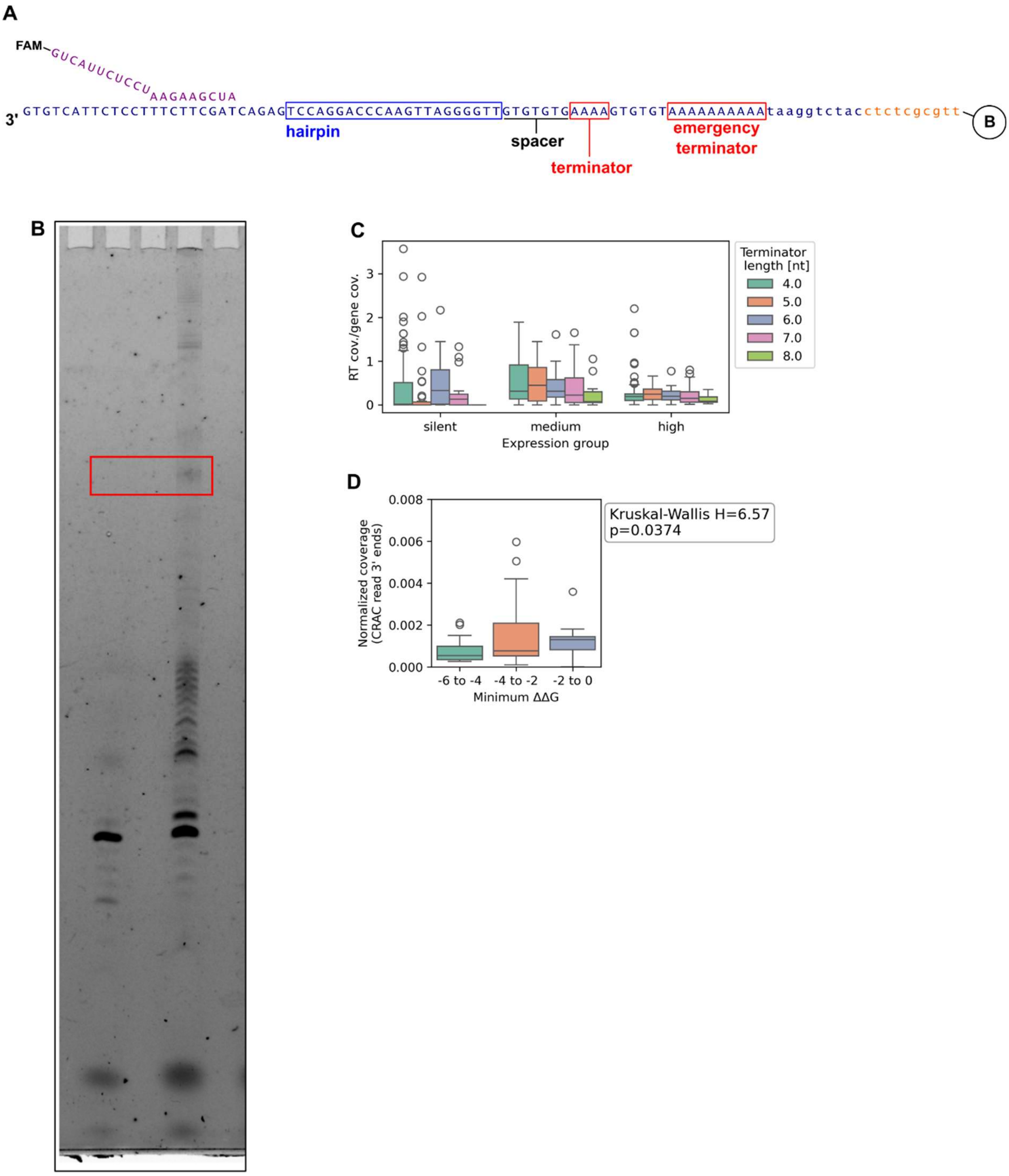

**Figure S7. Terminators have an impact on nascent transcript release by RNAPIII, related to Figures 5 and 6)**
A: Example sequence of biotinylated scaffolds used to assess transcript release in *in vitro* elongation assays. B – 5' biotin.
B: Uncropped image of gel shown in Fig. 6C. Cropped region marked with red box.
C: Readthrough intensity depends on terminator length in medium-, but not high-expression tRNA genes. Expression groups are same as ones in Fig S2.
D: Thermodynamic stability of transcription bubbles at the terminator, assessed by calculating the free energy difference (ΔΔG) of the RNA:DNA hybrid versus the reannealed DNA duplex.
